# Lgr5+ Stem Cells Maintain Apex Position in Cell Hierarchy of the Intestinal Epithelium During Homeostasis and Injury

**DOI:** 10.1101/2025.11.09.687498

**Authors:** Julian Chua, Lucy Driver, Masahiro Narimatsu, Mardi Fink, Lixia Luo, Jordan Tran, Arshdeep Kaur, Rida Habib, Jatin Roper, David G. Kirsch, Jeffrey L. Wrana, Chang-Lung Lee, Arshad Ayyaz

## Abstract

The cellular origin of intestinal epithelial homeostasis and regeneration has been a subject of continued debate, with recent models challenging the primacy of WNT-dependent Lgr5⁺ crypt base columnar (CBC) cells as the central intestinal stem cell population. Here, we revisit this question through quantitative integration of single-cell transcriptomic, chromatin accessibility, spatial, and lineage-tracing analyses across the proximal-to-distal axis of the small intestinal epithelium. Our data show that under homeostatic conditions, Lgr5⁺ cells exclusively sustain epithelial self-renewal in nearly all crypt–villus units along the entire length of the small intestine, a process for which R-spondin is indispensable. Following irradiation or chemotoxic injury, surviving Lgr5⁺ cells and their progeny reprogram into transient fetal-like cell states that initiate epithelial repair. Crucially, successful regeneration depends on the reactivation of canonical WNT/β-catenin signaling, as evidenced by increased TCF motif accessibility and upregulation of WNT target genes in newly forming Lgr5^+^ stem cells. Accordingly, pharmacological inhibition of WNT signaling blocks the reconstitution of Lgr5⁺ cells and crypt regeneration, leading to epithelial collapse. These findings reconcile prior controversies by demonstrating the central role of Lgr5⁺ CBC cells in epithelial self-renewal and regeneration following injury.

## INTRODUCTION

The intestinal epithelium is among the most rapidly self-renewing tissues in adult mammals, undergoing almost complete turnover every 3–5 days^1^. In the classical model, the constant replenishment of intestinal epithelial cells is sustained by crypt base columnar (CBC) cells, a population of actively cycling stem cells marked by Lgr5 expression and localized at the base of intestinal crypt epithelium^2–5^. In this model, Lgr5⁺ CBCs rely on continuous WNT/β-catenin signaling for their maintenance and proliferative capacity, giving rise to transit-amplifying (TA) cells that rapidly divide and differentiate into all mature epithelial lineages of the gut. This hierarchical cellular organization has served as the foundational model for understanding intestinal epithelial homeostasis^2,3,6^.

However, several subsequent studies have challenged the exclusivity and necessity of Lgr5⁺ CBCs in epithelial maintenance. In various injury or genetically modified contexts, alternative regenerative mechanisms have been proposed, including the activation of so-called “reserve” stem cells^7–10^, dedifferentiation of mature lineages^11–17^, and, more recently, a putative Fgfbp1⁺ isthmus progenitor cell (IPC) population located above the crypt base that may precede and give rise to Lgr5⁺ CBCs^18,19^. Reprogramming of surviving epithelial cells, induction of a fetal-like epitranscriptional program, and revival stem cell (revSC)-mediated reconstitution of fresh Lgr5^+^ cells have also been shown to regenerate the intestinal epithelium following somatic injury^20–23^.

One major complication in quantifying the contribution of Lgr5⁺ stem cells to intestinal regeneration is the widespread use of an earlier version of the *Lgr5-EGFP-IRES-CreERT2* reporter mouse line, in which the knock-in strategy disrupts endogenous Lgr5 function^2^. This results in a heterozygous Lgr5 knockout, mosaic Cre recombinase activity, and potentially incomplete epithelial labeling, which can lead to underestimation of the true contribution of Lgr5⁺ cells and misinterpretation that Lgr5⁻ populations may independently sustain epithelial turnover.

In addition, stem cell identity may vary along the length of the small intestine: although classically divided into three anatomical regions namely duodenum, jejunum, and ileum, recent studies have defined at least five discrete metabolic domains along the proximal-to-distal axis^24^. These observations raise the question of whether the stem cell role of Lgr5⁺ cells is spatially uniform or region-dependent, a question that has yet to be addressed quantitatively^24^.

Here, we revisit the question of cell and regenerative hierarchy along the entire length of the intestinal epithelium using a quantitative approach based on an *Lgr5-CreERT2* allele that preserves endogenous Lgr5 function^25^. We integrate this lineage tracing system with single-cell transcriptomics, chromatin accessibility profiling, spatially resolved gene expression analysis, and functional regeneration assays. Our findings show that nearly 100% of the crypt–villus units across the entire small intestine are maintained by Lgr5⁺ CBCs, including the turnover of Fgfbp1⁺ cells, which mainly mark the proliferative TA zone. Following severe injury, including irradiation and chemical injury, we find that Lgr5⁺ CBCs and their progeny undergo transient reprogramming into two previously described fetal-like states: a proliferative fetal-like CBC-like cell (FCC) and a quiescent revSC^20,26^. We further demonstrate that regeneration is contingent upon reactivation of WNT/β-catenin signalling, marked by increased chromatin accessibility at TCF-binding motifs. Pharmacological inhibition of WNT signalling impairs the emergence of revSC-derived fresh Lgr5^+^ stem cells and abolishes crypt regeneration, underscoring the essential role of this pathway in re-establishing stem cell identity and restoring normal epithelial turnover. Together, our findings reaffirm Lgr5⁺ CBCs as the central stem cell population governing both homeostatic renewal and injury-induced regeneration of the intestinal epithelium and define the transient fetal-like states that enable tissue repair upon somatic injury by re-instating the Lgr5^+^ stem cell pool.

## RESULTS

### Differential expression of Lgr5 and Fgfbp1 segregates stem cells and progenitor states during homeostasis

We integrated five independently generated single-cell transcriptomes of the mouse small intestine and first focused on the cellular composition of the intestinal epithelium under homeostatic conditions^6,20,22,24,26^. This integrated dataset, referred to as the ‘reference’, contained all major epithelial cell types, including Transit Amplifying (TA) cells, Enterocytes (EC), Goblet Cells (GC), Paneth Cells (PC), Tuft Cells (TC), Enteroendocrine Cells (EEC), Enterocyte progenitors (EC-prog), and Paneth-Goblet progenitors (PG-prog) **(Fig. 1a,b and Extended Data Fig. 1a,b)**. In addition, two populations of Lgr5^+^ CBCs were also observed, distinguished by their proliferative status: CBC1, representing relatively quiescent cells, and CBC2, representing proliferative cells **(Fig. 1c and Extended Data Fig. 2a,b)**. Two fetal-like cell populations were also detected in the reference dataset: quiescent revival stem cells (revSCs) and proliferative fetal-like CBC-like cells (FCCs), both of which are induced by injury^20,22,26^. Consistently, their abundance under homeostatic conditions was negligible, with FCCs comprising only 0.05% and revSCs 0.01% of the total epithelial population **(Fig. 1a,b and Extended Data Fig. 1a)**.

**Fig. 1:**
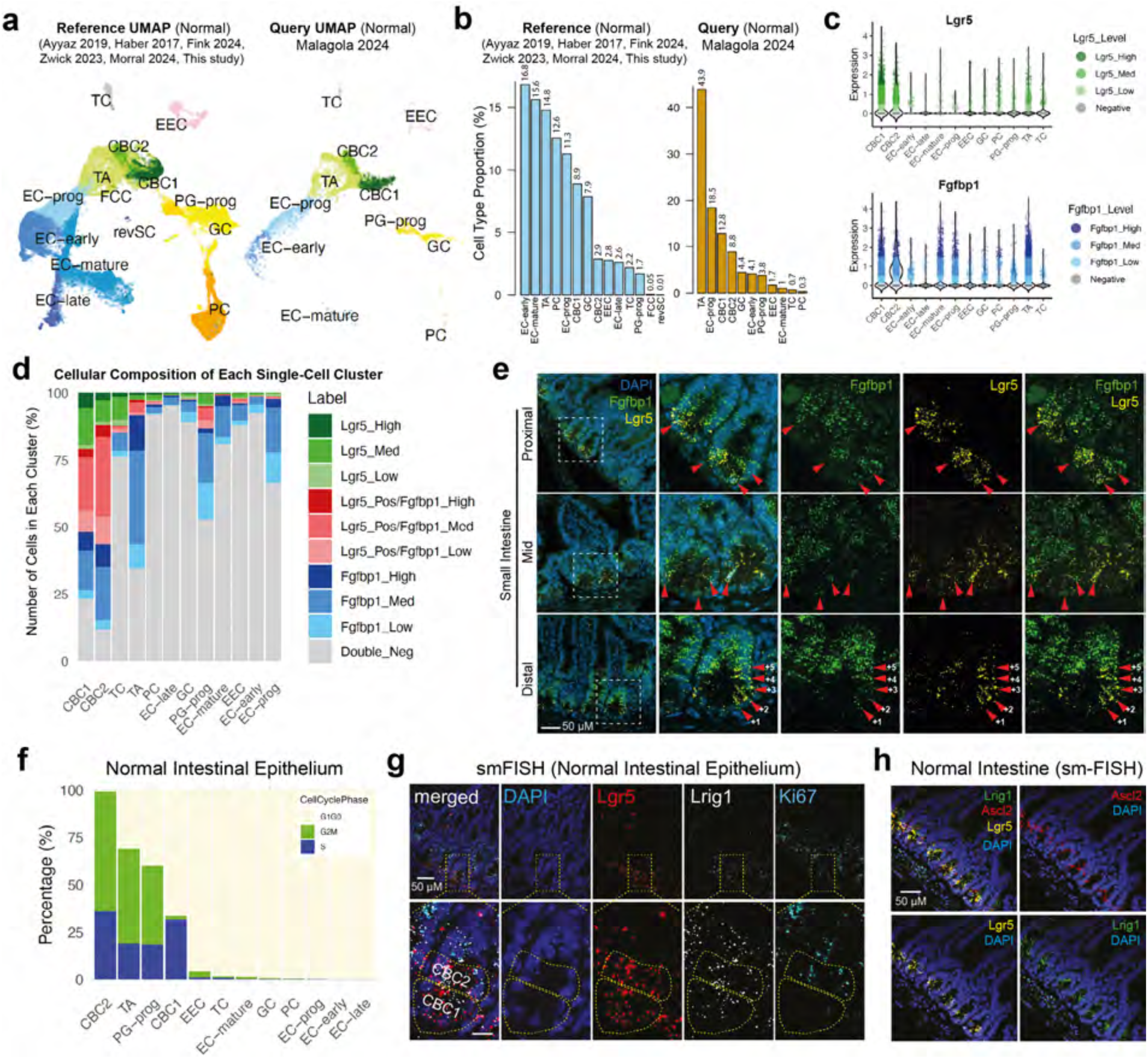
Integration and comparative analysis of homeostatic intestinal epithelial scRNA-seq datasets **(a)** UMAP reference map constructed by integrating six independent scRNA-seq datasets from whole intestinal epithelium under homeostatic conditions (right). Query scRNA-seq dataset of a whole intestinal epithelium under homeostatic conditions projected onto the reference UMAP (right) **(b)** Bar plot comparing cell percentage composition of major epithelial cell types between reference and query datasets. **(c)** Violin plots showing the average log2 fold-change expression of Lgr5 and Fgfbp1 across single cell types. Color intensity indicates expression levels (Negative: 0, Low: >0–0.5, Medium: 0.5–1.5, and High: >1.5), with darker colors representing higher expression. Lgr5 expression is depicted in shades of green, and Fgfbp1 expression is shown in shades of blue. **(d)** Stacked bar plot showing the relative proportion of epithelial cells within each annotated single-cell cluster, classified into mutually exclusive categories based on binned expression levels of Lgr5 and Fgfbp1. Log-normalized transcript levels were categorized as Low (>0–0.5), Medium (0.5–1.5), or High (>1.5). Cells expressing Lgr5 alone were grouped as Lgr5_Low, Lgr5_Med, or Lgr5_High (shades of green), and those expressing Fgfbp1 alone as Fgfbp1_Low, Fgfbp1_Med, or Fgfbp1_High (shades of blue). Cells co-expressing Lgr5 (>0) and Fgfbp1 at various levels were stratified as Lgr5_Pos/Fgfbp1_Low, Lgr5_Pos/Fgfbp1_Med, or Lgr5_Pos/Fgfbp1_High (shades of red). Cells negative for both genes were labeled as Double_Neg (grey). **(e)** Proximal, middle (mid), and distal small intestine sections were probed for Fgfbp1 and Lgr5 transcripts under homeostatic conditions. Fgfbp1 mRNA is predominantly localized in cells above the crypt base but is also co-expressed in Lgr5⁺ cells near the bottom of the crypts (red arrowheads). **(f)** Bar graph depicting the distribution of cell cycle phases (G1/G0, G2M, S) across all epithelial cell types in a normal mouse intestine. **(g)** sm-FISH staining of the whole normal mouse intestine (Top). sm-FISH staining of intact intestinal crypts, divided into spatial zones (1-3), probed for Lgr5, Lrig1, and Ki67 with DAPI counterstain. **(h)** sm-FISH staining for Lrig1, Ascl2, Lgr5, and DAPI of mouse crypts during homeostatic conditions.

We next compared the reference dataset to a recently published single-cell RNA-seq dataset, referred to as the ‘query’, which proposed the identification of “isthmus progenitor cells” (IPCs), that are marked by the expression of Fgfbp1, as the parent population of Lgr5⁺ CBC cells^18^. Similar cell types, including CBC1, CBC2, and TA cells were observed, although revSCs and FCCs were not detected in this dataset generated from normal intestines **(Fig. 1a,b)**. Due to their low or undetectable numbers in the normal intestine, we excluded revSC and FCC populations from further investigations of the homeostatic intestinal epithelium.

We next examined the distribution of *Lgr5* and *Fgfbp1* mRNA transcripts across intestinal epithelial cell types, which revealed that Lgr5 expression was predominantly restricted to CBC1 and CBC2 cell clusters, while Fgfbp1 transcripts exhibited broader distribution across additional cell types, including TA and early lineage-specific progenitors, such as EC-prog and PG-prog cells **(Fig. 1c and Extended Data Fig. 2b)**. These populations also expressed high levels of canonical cycle genes such as *Mki67*, *Pcna*, and *Top2a*, indicative of their proliferative state **(Extended Data Fig. 2c)**.

To evaluate the overlap in *Lgr5* and *Fgfbp1* expression across epithelial cell types, we classified individual cells into mutually exclusive categories based on binned transcript levels. Expression of each gene was stratified into Low, Medium, and High categories using log-normalized expression thresholds. Cells were then assigned to *Lgr5*-only (green), *Fgfbp1*-only (blue), or *Lgr5*⁺/*Fgfbp1*⁺ co-expressing groups (red), while cells lacking both transcripts were labeled as double-negative (grey), enabling a quantitative analysis of gene expression distributions across stem, progenitor and differentiated cell compartments **(Fig. 1d)**. In both CBC1 and CBC2 clusters, approximately 60% of cells expressed *Lgr5* (58% in CBC1 and 61% in CBC2), with the highest proportion of *Lgr5*_High (∼6.2%) and *Lgr5*_Med (∼13.7%) cells found in CBC1. While a substantial subset of CBC cells also co-expressed *Fgfbp1*, the nature of this co-expression differed markedly between the two CBC clusters. In CBC2, ∼44% of cells were double-positive for *Lgr5* and *Fgfbp1*, including ∼29.6% in the Lgr5_Pos/Fgfbp1_Med category, ∼10.1% in Lgr5_Pos/Fgfbp1_Low, and ∼4.3% in Lgr5_Pos/Fgfbp1_High. However, only ∼30.8% of CBC1 cells were double-positive, with the majority (∼20%) falling into the Lgr5_Pos/Fgfbp1_Med category. In stark contrast, TA cells exhibited the highest levels of *Fgfbp1* expression, with ∼57.6% of cells expressing *Fgfbp1* alone with minimal *Lgr5* expression. Among these, ∼34.7% fell into the Fgfbp1_Med category, underscoring the strong association of *Fgfbp1* with proliferative progenitor identity. Similar Fgfbp1-dominant profiles were seen in other proliferative compartments, such as EC-prog (∼31%) and PG-prog (∼33.7%), where Lgr5 positivity was negligible.

Taken together, these data delineate distinct epithelial programs, whereby Fgfbp1 marks actively cycling, progenitor-like populations, while Lgr5 marks stem-like compartments with variable proliferative potential, including subsets that co-express Fgfbp1.

### Heterogeneity within Lgr5⁺ CBC compartment is shaped by their position in crypts

To spatially resolve the CBC1 and CBC2 compartments in relation to Fgfbp1 gene expression pattern, we next performed RNAscope-based single-molecule in situ hybridization (sm-FISH). To assess regional differences, samples were further subdivided into proximal, middle (mid), and distal segments of the mouse small intestine. Consistent with our single-cell transcriptome analysis, and as described previously^19^, strong *Fgfbp1* expression was observed in the TA zone and extended toward the crypt–villus junction and into the base of the villi **(Fig. 1e)**. However, *Fgfbp1* expression was also readily detected in Lgr5^+^ cells in the crypt base with high *Fgfbp1* expression most prominent around the +3 to +5 cell positions and occasional *Fgfbp1* transcripts detected in Lgr5⁺ cells located at the bottom of the crypt base.

Next, we investigated the cell cycle dynamics across intestinal epithelial cell types. First, we examined the single-cell transcriptomes by assigning cell cycle stage scores to individual cells using previously established transcriptional signatures^2^. This approach enabled the identification of cells in the S phase, G2 or M phase (G2M phase), and G1 and/or G0 phase (G1G0 phase) **(Fig. 1f and Supplementary Table 1)**. The analysis spanned all epithelial cell types, revealing that CBC2, TA cells, and PG-prog populations were actively cycling. Notably, CBC2 possessed almost undetectable G1, suggesting rapid cycling, whereas CBC1 cells exhibited slower proliferation that was characterized by an extended S phase and almost undetectable G2/M. Most other epithelial cell types, including differentiated populations, remained largely quiescent.

To further delineate whether the Lgr5^+^ CBC compartment could be spatially split based on rate of proliferation, we employed sm-FISH. This analysis demonstrated that quiescent Lgr5^+^ Ki67^low/-^ cells, indicative of the CBC1 population, were localized predominantly at the crypt base **(Fig. 1g)**. In contrast, proliferative Lgr5^+^ Ki67^+^ cells, indicative of CBC2, were observed above the base, while cells in the upper crypt lacked *Lgr5* expression but were *Ki67*^+^, identifying them as TA cells. Interestingly, variable levels of *Lrig1* expression, another previously described marker of intestinal stem cells^8^, extended across all three crypt areas. Co-probing for *Lrig1*, *Ascl2* (yet another CBC marker) and *Lgr5* confirmed that *Lgr5* and *Ascl2* mRNA were colocalized in the crypt base and lower-to-middle crypt regions, whereas *Lrig1* expression extended further into the upper crypt areas and was evident at crypt-villus junctions **(Fig. 1h)**.

Together, these findings establish a spatial hierarchy within the Lgr5^+^ CBC compartment, comprising quiescent CBC1 cells at the crypt base and proliferative CBC2 cells positioned above them, followed by proliferative non-CBC Fgfbp1^+^ cells located in the upper crypt TA zone.

### Lgr5^+^ cells drive epithelial turnover throughout the intestine under homeostasis

It was recently claimed that Fgfbp1^+^ cells, but not Lgr5^+^ cells, possess an apical position in the intestinal epithelial hierarchy under homeostatic conditions. To test this, we first applied RNA velocity analysis using the reference dataset to infer lineage trajectories^27^. Velocity vectors were projected onto previously defined scRNA-seq clusters **(Fig. 1a)**. The directionality of lineage flow revealed a clear transition from CBC1 to CBC2 and subsequently to TA cells, which corresponds to their physical organization in the crypt **(Fig. 2a)**, and is consistent with a model in which Lgr5^+^ cells function as the principal self-renewing stem cell population. Notably, TA cells—enriched in *Fgfbp1*—did not exhibit reverse trajectories toward the CBC compartment but rather appeared as downstream progeny of CBC cells giving rise to differentiated lineages, reaffirming the canonical view of TA cells as short-lived transit progenitors.

**Fig. 2:**
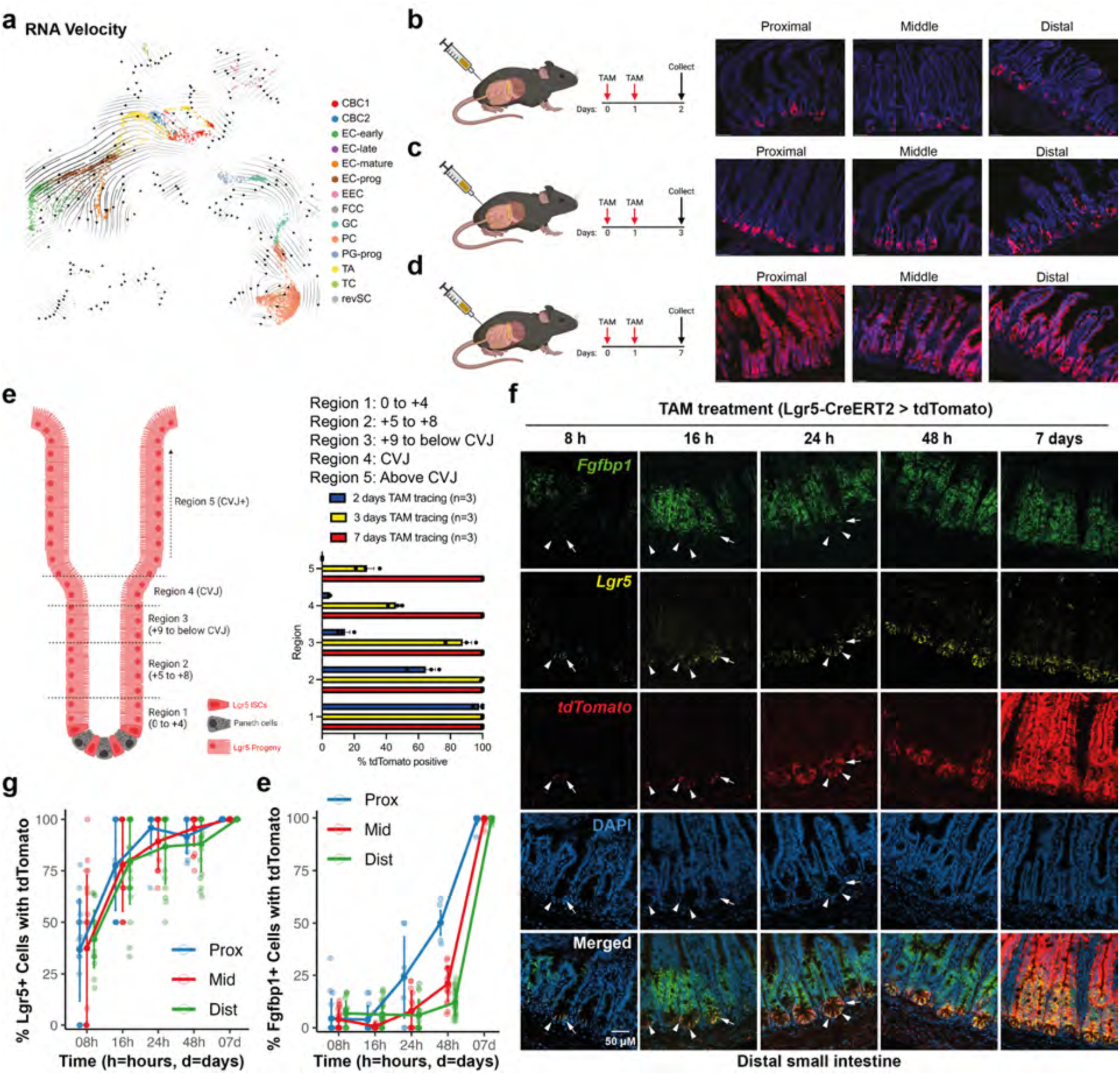
Lineage tracing reveals rapid labeling dynamics and spatial expansion of Lgr5+ cells in the intestinal epithelium **(a)** Velocity projected embeddings of murine full intestine at homeostasis. Velocity embeddings and velocity vectors (arrows) were calculated individually initially and then were projected onto UMAP embeddings of the scRNA reference map. **(b-d)** Schematic depicting unirradiated Lgr5-CreERT2; Rosa26-LSL-tdTomato mice given two injections of tamoxifen (TAM) and progeny labeled for 2 days (a), 3days (c), or 7 days (d) prior to harvest (day 0) of small intestines. Representative 20X images per segment per TAM dosing scheme. Scale bar 50um. Schematics created with BioRender^51^. **(e)** Schematic depicting zones 1-5 for quantification of Lgr5-tdTomato+ cells along with quantification of zones per TAM administration. n=3 mice per group. Red color indicates Lgr5+ stem cells/progeny, black indicates Paneth cells (intestinal cell types simplified for schematic purposes). Schematic created with BioRender^51^. **(f)** RNAscope in situ hybridization with Fgfbp1, Lgr5, and tdTomato probes, counterstained with DAPI, on distal small intestine sections treated with TAM for 8 hours to 7 days. TdTomatotdTomato⁺ clones emerge at 16 and 24 hours in single Lgr5⁺ cells at the crypt base. TdTomatotdTomato⁺Lgr5⁺Fgfbp1^low/^⁻ cells (white arrowheads) are observed at the crypt base, while tdTomato⁺Lgr5⁺Fgfbp1⁺ cells (white arrows) appear approximately +4 positions from the crypt base. **(g)** A line plot showing the percentage of Lgr5+ and tdTomato+ double positive cells across proximal, mid, and distal regions over time. The majority of Lgr5+ cells become tdTomato+ within 24h hours after Tamoxifen administration. **(e)** A line plot showing the percentage of Fgfbp1+ and tdTomato+ double positive cells across proximal, mid, and distal regions over time. tdTomato labeling increases progressively over time, with nearly all Fgfbp1⁺ cells labeled by 7 days.

Next, we wanted to experimentally assess the contribution of Lgr5⁺ cells by employing lineage tracing, a technique where previous studies used *Lgr5-EGFP-IRES-CreERT2* mice, in which an *IRES-CreERT2* cassette was inserted at exon 1 of the *Lgr5* gene^25^. This resulted in functional heterozygosity, as homozygous knockout of *Lgr5* is embryonic lethal. Furthermore, since Lgr5 itself is important for R-spondin-mediated promotion of WNT signalling via inhibition of Znfr3 and RNF43, this may have an impact on WNT pathway activity. In addition, these mice exhibit mosaic expression of the transgene in the intestinal epithelium, which may confound lineage-tracing results and lead to an underestimation of the actual contribution of Lgr5⁺ CBCs to epithelial maintenance^2^.

Therefore, to ensure preservation of *Lgr5* gene function while investigating the contribution of Lgr5^+^ and Lgr5^-^ cells to epithelial turnover, we utilized a more recently developed *Lgr5-CreERT2* mouse line, in which an *IRES-CreERT2* cassette is inserted into the 3’ untranslated region (UTR) of the *Lgr5* locus, without disrupting *Lgr5* gene function^25^. We crossed these mice with *Rosa26-LSL-tdTomato* reporter mice and initiated lineage tracing under homeostatic conditions using two doses of TAM administered on consecutive days. Intestines were subsequently harvested at 2-, 3-or 7-days post-TAM treatment initiation **(Fig. 2b-d and Extended Data Fig. 3a-d)**. This showed that at day 2, nearly 100% of crypt bases (region1; position 0 to +4) possessed tdTomato^+^ flask-shaped Lgr5^+^ CBCs, whereas Paneth cells were tdTomato^-^.

Labelling in region 2 was less prevalent (∼60% of cells) (mid to upper TA and CVJ), with no cells labeled in region 5 (villus) **(Fig. 2b,e and Extended Data Fig. 3a,b)**. At day 3, extensive labelling of cells now extended into region 2 (∼100%; position +5 to +8) and region 3 (∼88%; position +9 to below CVJ) and was 44% in region 4, while the villi (region 5) now contained ∼28% tdTomato+ cells **(Fig. 2c,e and Extended Data Fig. 3a,c)**. After 7 days of pulse, nearly 100% of intestinal epithelial cells throughout the crypt-villus axis were labeled by tdTomato except for long-lived Paneth cells **(Fig. 2d,e and Extended Data Fig. 3a,d)**. These results demonstrate that the entire epithelium of the murine small intestines, except for Paneth cells, is repopulated by the progeny of Lgr5^+^ cells within 7 days at homeostasis.

These results contradict previous claims that Lgr5⁺ cells can be repopulated by Fgfbp1⁺ cells within 7–14 days^19^. Therefore, to quantify Lgr5⁺ lineage-tracing kinetics relative to Fgfbp1⁺ cells, we performed temporally resolved sm-FISH targeting *Lgr5*, *Fgfbp1*, and *tdTomato* transcripts across proximal, middle, and distal regions of the small intestine (**Fig. 2f** and **Extended Data Fig. 4a,b**). To quantify, we used Cell Profiler to first create single cell masks, and counted *Lgr5*, *Fgfbp1* and *tdTomato* transcripts in single cells^28^ (**Extended Data Fig. 4c**). This analysis showed that, on average, 21%, 16%, and 36% of Fgfbp1⁺ cells in the proximal, middle, and distal regions of the small-intestinal epithelium, respectively, were also Lgr5⁺ (**Fig. 2f** and **Extended Data Fig. 4d**).

Next, we measured the fraction of Lgr5⁺ cells that were tdTomato-labeled across proximal, middle, and distal small-intestine regions over time (**Fig. 2g**). A substantial ∼36–40% of Lgr5⁺ cells were already tdTomato⁺ at 8 h (Proximal 36.2%, Middle 36.8%, Distal 40.1%). This increased to ∼67–70% by 16 h (Proximal 67.5%, Middle 70.4%, Distal 69.0%) and reached ∼79–86% by 24 h (Proximal 85.7%, Middle 82.5%, Distal 78.7%), at which point the majority of crypt-resident Lgr5⁺ cells were labeled, with the residual unlabeled Lgr5⁺ cells observed higher along the villus axis and a few in the non-epithelial lamina propria, as previously described^29^. Labeling rose further to ∼84–92% at 48 h (Proximal 87.8%, Middle 91.9%, Distal 83.6%) and 99% by 7 days (Proximal 100%, Middle 97.6%, Distal 99.5%).

Consistent with transcriptional overlap between Lgr5^+^ and Fgfbp1^+^ cells, a small fraction of Fgfbp1⁺ cells were labeled at the earliest time points (**Fig. 2e**). At 8 h, ∼6–12% of Fgfbp1⁺ cells were tdTomato⁺ (Proximal 11.7%, Middle 6.5%, Distal 8.4%), and this proportion remained low at 16 h (Proximal 10.1%, Middle 6.6%, Distal 9.7%). By 24 h, labeling had increased, particularly in the proximal region (Proximal 39.3%, Middle 14.0%, Distal 8.5%), and rose further by 48 h (Proximal 53.7%, Middle 19.7%, Distal 13.9%). By 7 days, nearly all Fgfbp1⁺ cells across the small intestinal epithelium were tdTomato⁺ (Proximal 93.8%, Middle 98.1%, Distal 98.6%). Given that tdTomato labeling is a permanent mark of Lgr5⁺ cells and their progeny, the progressive increase in tdTomato⁺ Fgfbp1⁺ cells over time demonstrates that the Fgfbp1⁺ population arises from Lgr5⁺ stem cells within 7 days.

### R-spondin is indispensable for intestinal epithelial turnover

Our lineage tracing demonstrates that Lgr5⁺—but not Fgfbp1⁺—cells occupy the apex of the epithelial hierarchy along the entire length of the small intestine, and that Fgfbp1⁺ cells are instead direct progeny of Lgr5⁺ cells. These findings not only contradict the notion that Fgfbp1⁺ cells are precursors of Lgr5⁺ cells but also cast doubt on prior claims that epithelial turnover is R-spondin–independent, a conclusion based on the asserted dispensability of R-spondin for Fgfbp1⁺ cell self-renewal^29^. Because R-spondin is important for maintaining WNT signaling ^2,3^, we performed a comprehensive molecular analysis of WNT signaling activity, beginning with differential gene expression across immature epithelial populations (CBC1, CBC2, TA, PG-prog, and EC-prog) **(Supplementary Table 2)**. This analysis revealed distinct gene expression patterns. For instance, conventional β-catenin target genes such as Lgr5, Olfm4, Axin2, and Ascl2 were enriched in both CBC1 and CBC2 populations, as were Wnt receptor genes like Fzd1, Fzd6, and Lrp5. However, other WNT receptors, including Fzd4, Fzd8, Fzd9, and Lrp6, showed significant expression in TA, PG-prog, and EC-prog populations **(Extended Data Fig. 5a)**. Notably, Fzd5 expression extended to non-CBC cell types, similar to that of Lrig1, a finding further validated by immunofluorescence staining of Lgr5 (GFP), Lrig1, and Fzd5 **(Extended Data Fig. 5b)**. These observations indicate that WNT/β-catenin signaling is not strictly confined to CBCs but may play a broader role in influencing other cell populations within the intestinal epithelium, including differentiation of progenitors.

To test this hypothesis, we next evaluated WNT signaling activity using a β-catenin gene signature^30^. This revealed elevated β-catenin activity in CBC2, TA cells, and PG-prog cells, but surprisingly, not in CBC1 and EC-prog cells **(Extended Data Fig. 5c** and **Supplementary Table 3)**. The differential activity of WNT signalling in distinct crypt cell populations prompted us to test the role of the WNT pathway in organoids that model intestinal dynamics.

Canonical WNT signaling relies on R-spondin ligands, which bind to Lgr5 and inhibit the ubiquitin ligases Znrf3 and Rnf43, thus preventing the degradation of WNT receptors and sustaining β-catenin activity^31^. Both Znrf3 and Rnf43 were differentially expressed in CBC1, CBC2, TA cells, and PG-prog cells, underscoring their potential role in modulating WNT activity in these populations **(Extended Data Fig. 5c)**. To directly test the functional role of R-spondin, we established intestinal organoids from C57BL/6 mice. When cultured without recombinant R-spondin1 (Rspo1) for 3 days, these organoids showed complete loss of crypt formation and proliferation, as confirmed by Ki67 immunostaining, whereas organoids cultured in Rspo1 contained Ki67+ crypt structures **(Extended Data Fig. 5d)**. Repeating these experiments using Lgr5-GFP reporter organoids demonstrated a similar outcome: the withdrawal of Rspo1 abolished Lgr5^+^ CBCs and Paneth cell formation, the latter identified by Lyz1 staining **(Extended Data Fig. 5e)**. Further validation by staining for Olfm4, another CBC marker, revealed a complete absence of Olfm4^+^ cells in the absence of Rspo1 **(Extended Data Fig. 5f)**.

Next, organoids cultured for 3 days in Rspo1-deficient medium were either returned to Rspo1-supplemented medium for 2 days or maintained without Rspo1 for a total of 5 days; parallel controls were continuously cultured with Rspo1 for 5 days (**Extended Data Fig. 5g**). Regardless of reintroduction, organoids subjected to the initial 3-day withdrawal underwent progressive degeneration and collapsed by day 5, whereas normal organoid growth was observed upon continuous Rspo1 treatment. These data indicate that even transient Rspo1 deprivation triggers an irreversible loss of crypt stem-cell–dependent regenerative capacity. Similar studies in vivo point to stromal Rspo3 as an important promoter of WNT signalling in the crypt^32^.

Together, these findings demonstrate that R-spondin is not only essential for maintaining the Lgr5⁺ CBC, TA and early progenitor populations but is also required for driving any proliferation in the intestinal epithelium as well as enabling differentiation of Paneth cells. These results thus highlight the indispensable role of R-spondin–mediated WNT/β-cateninsignaling in regulating epithelial cellular turnover in the intestine under steady-state conditions.

### Distinct proliferative and quiescent fetal-like programs are activated upon severe injury

Integrating single-cell transcriptome profiles of the intestine across multiple time points after irradiation, measured as days post-irradiation (dpi), allowed us to track cellular dynamics in real-time during intestinal epithelial regeneration. To this end, we calculated fold changes for individual cell types, which revealed marked compositional shifts across the epithelium, which were consistent with previous findings^20^ (**Fig. 3a-c**). In particular, the homeostatic stem-cell compartment was progressively depleted: CBC1 fell from 8.9% (0 (dpi) to 4.6% (1 dpi), 1.7% (2 dpi), and 0.063% (3 dpi), and CBC2 declined from 2.9% (0 dpi) to 3.3% (1 dpi), 1.6% (2 dpi), and 0.013% (3 dpi). Despite a minor uptick at 1 dpi for CBC2, both populations exhibited >99% loss by 3 dpi, indicating rapid loss of crypt-resident stem cell phenotypes following irradiation.

**Fig. 3:**
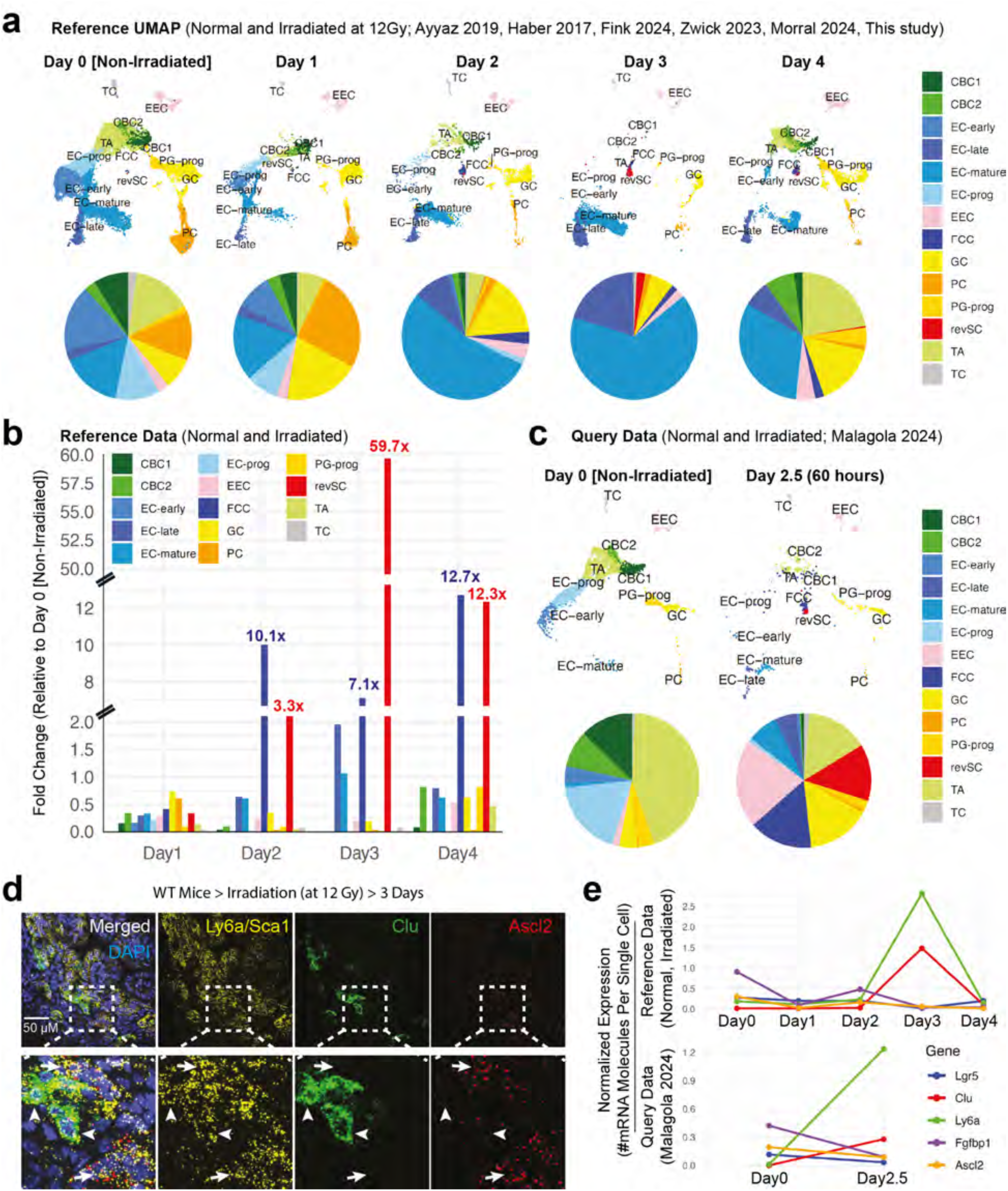
Cellular and molecular remodelling of the intestine epithelium after radiation-induced damage **(a)** Reference UMAP of the intestinal epithelium at homeostasis (day 0) and at 1-, 2-, 3-, and 4-days post-IR, annotated with pie charts indicating proportional cell type changes over time. **(b)** Bar graph quantifying fold-change in cell type abundance in the reference dataset relative to day 0. **(c)** UMAP visualization of query datasets from day 0- and 60-hours post-IR, with pie charts illustrating cell type shifts. **(d)** sm-FISH staining of the mouse intestine 3 days post-12 Gy IR stained with Ly6a/ Sca1, Clu, and Ascl2. **(e)** Line plots showing normalized RNA molecule counts per cell for key intestinal epithelial markers over the course of regeneration (day 0 to day 4) in both reference and query datasets.

Importantly, epithelial composition trended back toward homeostasis by 4 dpi, including the stem and transit compartments rebounding sharply: CBC1 rose from 0.063% at 3 dpi to 2.27% at 4 dpi and CBC2 from 0.013% to 7.59%, while TA recovered from 0.063% to 22.4%. Concurrently, revSC population, which has previously been shown to generate fresh Lgr5^+^ cells^20^ declined significantly, dropping from 2.27% at 3 dpi to 0.39% at 4 dpi, consistent with an ongoing but active return toward pre-injury architecture of the stem cell compartment. Query data from Malagola et al. collected at 2.5 dpi with 12 Gy showed a similar trend, albeit with substantially higher proportions of injury-induced revSC (13.6%) and FCC (15.2%) **(Fig. 3c)**. These differences may arise from variations in sample preparation, such as using fluorescence-activated cell sorting (FACS) to deplete B6A6+ villi cells from Epcam+ single cell suspensions before subjecting them to sc-RNA-seq^18,33^.

Interestingly, in a complete contrast with revSC population, proportion of FCCs rose from 1.53% at 3 dpi to 2.27% at 4 dpi, prompting us to further understand the distinctive nature of injury-associated populations by examining the expression of *Ly6a*, *Clu*, and *Ascl2*, which is a CBC marker and expressed sporadically in FCC, by sm-FISH as previously described^20^ (**Extended Data Fig. 6a**). At 3 dpi Ly6a+ cells could be readily partitioned into Clu+ versus Ascl2+ populations (**Fig. 3d**), thus demonstrating that fetal-like cells induced upon irradiation consist of two distinct populations: Ly6a^+^ Clu^-^ Ascl2^+^ FCC and Ly6a^+^ Clu^+^ Ascl2^-^ revSC. Together, these results show that revSCs and FCCs are mutually exclusive injury-induced states with distinct kinetics: revSCs decline as homeostatic Lgr5⁺ stem cells re-emerge, whereas FCCs expand during this interval.

### Expression of Fgfbp1 is primarily associated with the homeostatic crypts

Our results identify revSC and FCC populations as the two prominent injury-associated transcriptional states that could potentially produce fresh Lgr5^+^ CBC cells and thereby regenerate the entire intestinal epithelium. An alternative model of epithelial repair posits that Fgfbp1⁺ cells replenish the Lgr5⁺ ISC pool. We found that while Fgfbp1 expression was predominantly associated with multiple populations in homeostatic crypts—especially TA, CBC1, and CBC2 cells—it declined during the reprogramming phase of regeneration, with only a few transcripts detectable in revSC and FCC populations, especially at their peak expansion stages at 3 dpi and 4 dpi, respectively (**Fig. 3a,b** and **Extended Data Fig. 6a,b**). To further confirm these trends, we performed pseudobulk analysis to assess regeneration-dependent regulation by quantifying the total mRNA transcripts of Fgfbp1 per cell at each time point and compared to the fetal-like gene signature marker Ly6a, the revSC marker Clu, and the CBC markers Lgr5 and Ascl2 **(Fig. 3e)**. This analysis revealed a significant suppression of Fgfbp1 similar to the loss of TA cells and CBC markers Lgr5, and Ascl2, whereas transcripts for Clu and Ly6a were markedly elevated at days 2.5 and 3, in both the reference and query datasets. Collectively, these data indicate that Fgfbp1 is not a feature of damage-induced regenerative cell states and is downregulated during crypt reprogramming, consistent with depletion of homeostatic TA and CBC compartments in the regenerating crypt.

### Revival stem cells transition through fetal-like CBC-like cells

The homeostatic stem cell populations, CBC1 and CBC2, continue to decline until 3 dpi (**Fig. 3a-c**). By 4 dpi, both populations start to expand for the first time post-irradiation, coinciding with a rapid decline in the revSC population, consistent with prior findings that de novo Lgr5⁺ cells are repopulated by revSCs^20^. Importantly, the decline in revSCs also coincides with an approximately two-fold expansion of the FCC population, prompting us to further investigate relationships between individual homeostatic stem and regenerative cell populations. For these analyses, we specifically focused on stem cell dynamics at 4 dpi and proceeded by performing de novo clustering and differential gene expression analyses comparing CBC1, CBC2, FCC, and revSCs. This revealed multiple overlapping and transitioning populations (**Fig. 4a**).

**Fig. 4:**
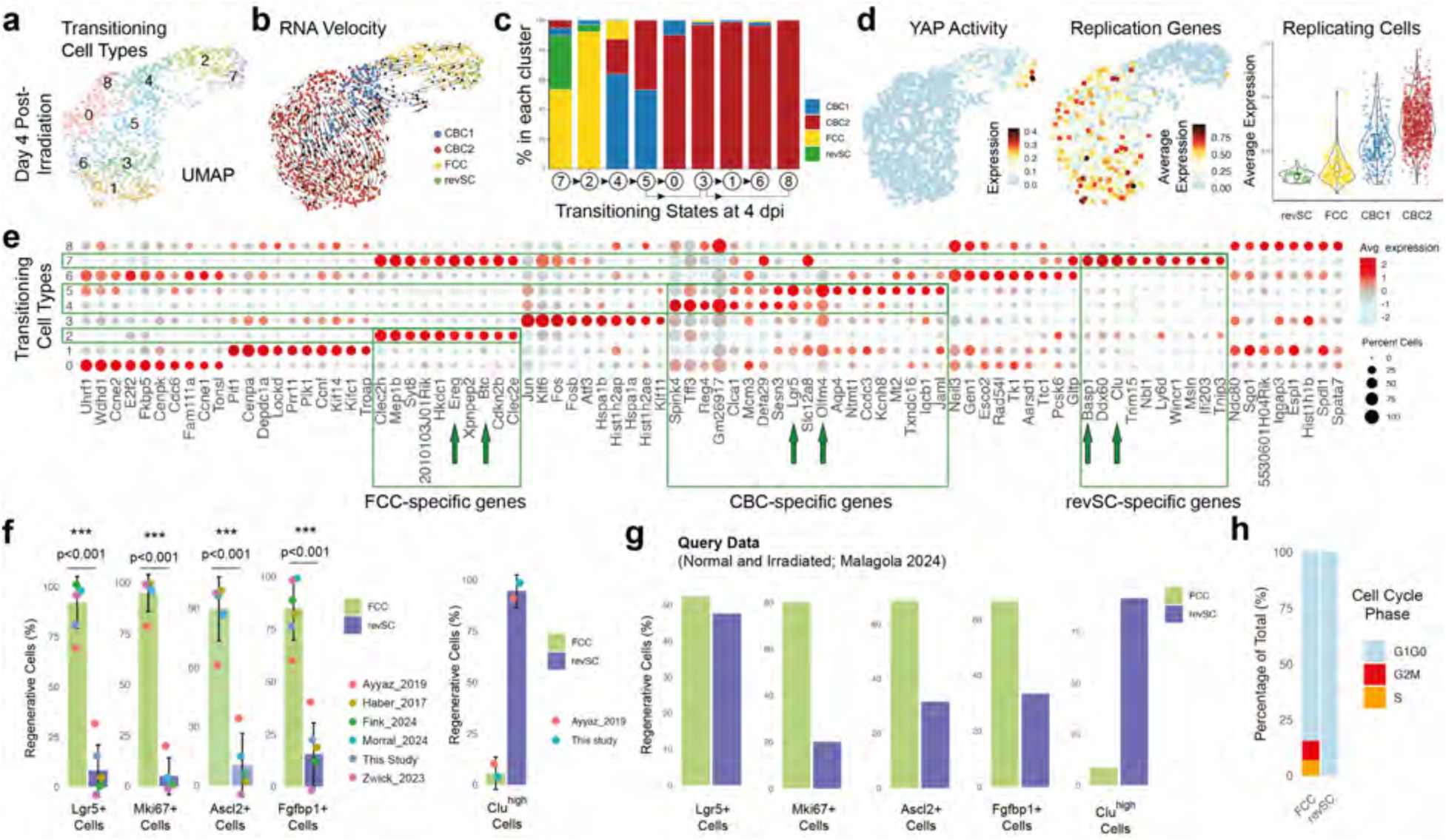
Characterization of regeneration associated epithelial subtypes following irradiation-induced injury **(a)** scRNA UMAP of CBC1, CBC2, FCC, and revSC compartment 4 days post-irradiation. **(b)** UMAP of transitioning cell types CBC1, CBC2, FCC, and revSCs with RNA velocity vectors (arrows) on original scRNA UMAP embeddings. **(c)** Bar graph depicting the percentage of CBC1, CBC2, FCC, and revSC cells contributing to each cluster transition state. **(d)** Feature plot of YAP activity (left), replicating gene signature (calculated as average expression of Mki67, Pcna, Top2a, Cdk1, Ccnb1, Birc5, Aurka, Mcm5 and Mcm6; middle) and a violin plot showing enrichment of replicating gene signature across regenerative (revSC and FCC) and homeostatic stem cell (CBC1 and CBC2) populations at 4 dpi (right). **(e)** Dotplot showing results from differential gene expression analysis of transitioning states at 4 dpi. **(f)** Classification of Sca1^+^ cells into FCCs and revSCs, with quantification of cells expressing regeneration-associated genes. Each dot represents and independent biological replicate from an independent scRNA-seq dataset. Significance was calculated using a two-tailed t-test. **(g)** A bar plot of the percentage of Lgr5+, Mki67+, Ascl2+, Fgfbp1+, and Clu^high^ Cells in FCC (Green) and revSC (Blue) in the query dataset. **(h)** Bar graph showing cell cycle phase distribution (G1/G0, S, G2/M) in FCCs and revSCs.

Concordant trajectory inference using RNA velocity and Slingshot delineated a regenerative axis in which cluster #7—co-enriched for FCCs and revSCs—lies upstream of the expanding FCC-dominant cluster #2. From this node, lineage trajectories progress to clusters #4 and #5, both predominantly enriched for CBC1, which subsequently give rise to the remaining five CBC2-enriched clusters^27,34^ (**Fig. 4b,c** and **Extended Data Fig. 6c,d)**. This conversion of regenerative states into homeostatic states was accompanied by reduced Yap activity: highest in revSCs, moderate to low in FCCs, and almost none in CBC populations (**Fig. 4d**).

Conversely, expression of a replicative gene signature, which was suppressed in revSCs, was progressively induced in FCCs and the de novo CBC1 and CBC2 compartment2. Consistent with the association of Fgfbp1 with homeostatic crypts, *Fgfbp1* expression was also sporadically induced in the CBC1 and CBC2 compartments (**Extended Data Fig. 6e**).

Our trajectory analysis placed the FCC population at the junction between revSC and CBC compartments, prompting us to examine its association with both. First, we performed comprehensive differential gene-expression profiling of transitioning phenotypes. Consistent with the confinement of most revSCs to cluster #7, canonical revSC genes, such as Clu and Basp1, were predominantly expressed in cluster #7, whereas several FCC-associated genes, including Ereg and Btc, were expressed in both cluster #7 and the FCC-enriched cluster #2 (**Fig. 4e**). Because cluster #7 at 4 dpi contains both FCC and revSC cells in a transitional state, we performed a direct comparison by pooling both cell types across all time points, defining gene-positive cells as those with log2-normalized expression > 0, and comparing their distributions within FCC and revSC clusters across all datasets and time points (**Fig. 4f**). This analysis showed that expression of CBC markers Lgr5, Ascl2, cell-cycle genes (e.g., Mki67), and the proposed “IPC” marker Fgfbp1 was largely restricted to FCC cells, whereas Clu-expressing cells were predominantly found in the revSC population. Similar results were obtained in the query dataset from Malagola et al. (**Fig. 4g**). Furthermore, cell-cycle analysis revealed that 15.6% of FCC cells were in S or G2/M phases based on a phase-specific gene signature, whereas 100% of revSCs were in G1/G0 (**Fig. 4h**). Taken together, these results identify the FCC population as a transitional state through which quiescent revSCs reconstitute the proliferative homeostatic stem-cell compartment.

### Lgr5^+^ cells and their progeny repopulate the damaged intestinal epithelium following different severities of radiation injury

Ionizing radiation causes acute injury to the small intestinal epithelium in a dose-response manner^35^. In general, radiation doses at 12 Gy and above are considered high doses that cause severe damage to the small intestinal epithelium of mice^35,36^. For example, we have shown that in C57BL/6 mice, 12 Gy and 14 Gy sub-total-body irradiation (SBI) deplete >80% and >95% of proliferating crypts in the small intestines 4 days after irradiation, respectively^37^. To determine the contribution of Lgr5^+^ cells and their progeny to intestinal regeneration after different severities of radiation injury, we injected *Lgr5-CreERT2; Rosa26-LSL-tdTomato* mice with TAM days 2 and 1 (TAM -2, -1d) before irradiation to label CBC and lower TA cells **(Fig. 2f and 5a)** or TAM at days 7 and 6 (TAM -7, -6d) before irradiation to label all epithelial cells along the crypt-villus axis except for Paneth cells **(Fig. 5b)**. These mice were then exposed to 0, 10, 12, or 14 Gy SBI, and the small intestines were harvested 10 days post-irradiation to assess lineage tracing events **(Fig. 5a-g and Extended Data Fig. 7a-i and Fig. 8a-i)**. Of note, while 10 and 12 Gy SBI did not cause lethality in *Lgr5-CreERT2; Rosa26-LSL-tdTomato* mice, 14 Gy SBI caused gastrointestinal acute syndrome (GI-ARS) in more than 50% of mice within 10 days after irradiation **(Extended Data Fig. 7a and Fig. 8a)**. Therefore, in the 14 Gy SBI group, only mice that survived 10 days after irradiation were analyzed.

**Fig. 5:**
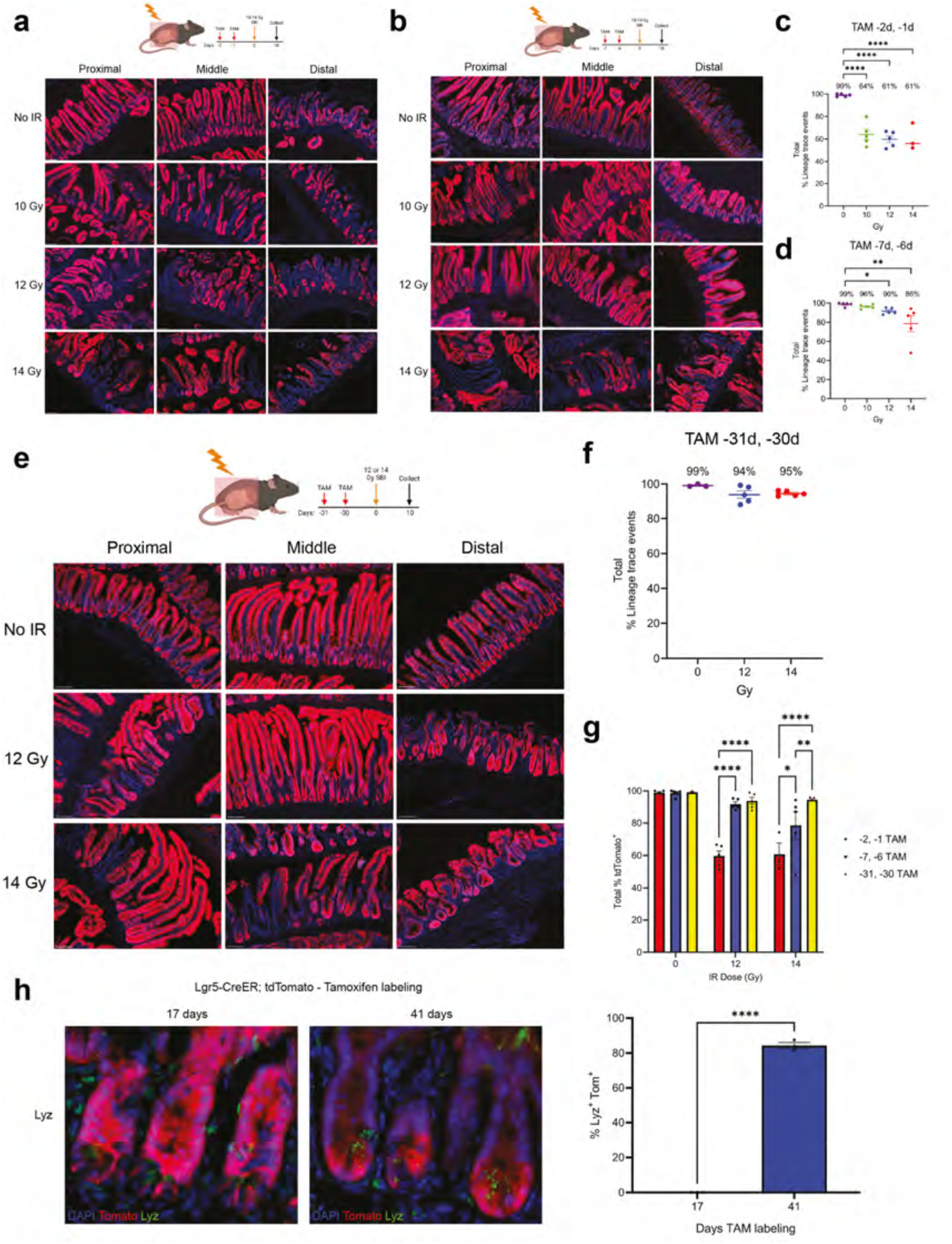
Lineage tracing reveals effects of irradiation on Lgr5+ stem cell progeny in the intestinal epithelium **(a-b)** Schematic depicting Lgr5-CreERT2; Rosa 26-LSL-tdTomato mice were given TAM to label Lgr5 progeny for 2 days (a) or TAM -7 days (b) prior to irradiation of either 0-14 Gy SBI and small intestines were harvested 10 days later. Representative 10X images per segment per irradiation dose 0-14 Gy SBI. Scale bar 100um^51^. **(c)** Quantification of 2 day pre-IR labeling total lineage trace events p<0.0001 No IR vs. 10, 12, and 14 Gy SBI. Each dot represents one mouse. Statistics performed per nonparametric one-way ANOVA. **(d)** Quantification of 7 day pre-IR labeling total lineage trace events p=0.04 No IR vs. 12 Gy SBI; p=0.001 No IR vs. 14 Gy SBI. Each dot represents one mouse. Statistics performed via nonparametric one-way ANOVA. Schematic created with BioRender. **(e)** Schematic depicting Lgr5-CreERT2; Rosa26-LSL-tdTomato mice given two injections TAM and Lgr5 progeny labeled for 31 days prior to irradiation of either 12 or 14 Gy SBI and small intestines harvested 10 days later. Representative 10X images per segment per irradiation dose. Scale bar 100um. Schematic created with BioRender^51^. **(f)** Quantification of 31 day pre-IR labeling total lineage trace events. Each dot represents one mouse. Statistics performed via 2way ANOVA. **(g)** Quantification of 2 day pre-IR, 7 day pre-IR, and 31 day pre-IR labeling, -30d total lineage trace events p=<0.0001 2 day pre-IR labelling 12 Gy SBI vs. 7 day pre-IR labeling 12 Gy SBI; p=<0.0001 2 day pre-IR lableing 12 Gy SBI vs. 31 day pre-IR labeling 12 Gy SBI; p=0.01 2 day pre-IR labeling 14 Gy SBI vs. 7 day pre-IR labeling 14 Gy SBI; p=<0.0001 2 day pre-IR labeling 14 Gy SBI vs. 31 day pre-IR labeling 14 Gy SBI; p=0.009 7 day pre-IR labeling 14 Gy SBI vs. 31 day pre-IR labeling 14 Gy SBI. Each dot represents one mouse. Statistics performed via nonparametric one-way ANOVA.

Quantification of lineage tracing events showed that labeling progeny for 2 days pre-IR and subsequent tissue harvest 10 days post-IR, 99% of crypts were labeled by tdTomato^+^ 10 days after sham irradiation (0 Gy), whereas in the irradiated group, only 64% of crypts were tdTomato^+^ after 10 Gy SBI, 61% of crypts were tdTomato^+^ after 12 Gy SBI, and 61% of crypts were tdTomato^+^ after 14 Gy SBI **(Fig. 5a,c)**. The percentage of tdTomato^+^ crypts was significantly lower in irradiated cohorts compared to unirradiated mice; however, we did not observe a clear dose response in tdTomato^+^ cells between mice that received 10, 12, and 14 Gy SBI **(Fig. 5a,c)**. Similar lineage tracing events were observed across the proximal, middle, and distal small intestines **(Extended Data Fig. 7c-i)**. In contrast to the results from the 2 day pre-IR tracing, upon labeling cells for 7 days prior to irradiation and subsequent 10 days later harvest, 99% of crypts were labeled by tdTomato^+^ 10 days after sham irradiation (0 Gy), compared to 96%, 90% and 86% after 10, 12 and 14 Gy SBI, respectively **(Fig. 5b,d)**. Notably, differences in lineage tracing events were not statistically significant between unirradiated and 10 Gy SBI cohorts, while the percentage of tdTomato^+^ crypts in either 12 Gy SBI or 14 Gy SBI cohorts was significantly lower compared to the unirradiated group (0 Gy) that was evident across the proximal, middle, and distal small intestines or 14 Gy-treated mice **(Extended Data Fig. 8c-i)**.

The more efficient lineage tracing upon treatment with TAM at -7, -6d is consistent with an apex position for Lgr5^+^ CBCs in the intestinal epithelium. However, 10 and 14% of regenerated crypts remained unlabelled after 12 and 14 Gy irradiation. This could be evidence for the existence of a progenitor upstream of Lgr5^+^ ISCs. Alternatively, a population of long-lived Lgr5^+^ offspring that are not labelled within 7 days before damage might be important for intestinal regeneration following severe injury induced by high doses of radiation exposure. This prompted us to consider if Paneth cells, which have a turnover rate of more than 30 days^38^, might contribute significantly to intestinal regeneration. To test this, we injected *Lgr5-CreERT2; Rosa26-LSL-tdTomato* mice with two doses of TAM to label all Lgr5 progeny for 31 days before 0, 12, and 14 Gy SBI and harvesting tissues 10 days later **(Fig. 5e and Extended Data Fig. 9a,b)**. Our results show that 99% of crypts were labeled by tdTomato^+^ 10 days after sham irradiation (0 Gy), and after 12 and 14 Gy SBI tdTomato+ crypts rose to 94% and 95% respectively **(Fig. 5e,f)**. The increased efficiency of labelling at 14 Gy SBI was significantly higher than the 7 day pre-IR tracing cohort **(Fig. 5g)**, and was observed across the entire proximal, middle, and distal axis of the small intestines **(Extended Data Fig. 9c-h)**. Staining of Lyz^+^ Paneth cells in unirradiated mice further revealed that while no Paneth cells were positive for tdTomato 17 days after TAM, approximately 80% of Paneth cells were labeled by tdTomato 41 days after TAM treatment **(Fig. 5h)**.

Together, the results from our lineage tracing experiments demonstrate that following 10 to 14 Gy SBI, more than 95% of the mouse intestinal epithelium is regenerated from Lgr5^+^ CBCs and their differentiated progeny. Of note, this regenerative response is affected by the severity of injury. While Lgr5^+^ cells and their early progeny labeled by TAM treatment 7 days pre-IR are sufficient to regenerate the entire irradiated epithelium of the small intestines after 10 Gy SBI. In mice exposed to 14 Gy SBI, which causes the most severe injury, Lgr5-derived Paneth cells make a significant contribution to regenerating the damaged intestinal epithelium.

### Epigenetic landscape of regenerating epithelium indicates re-activation of WNT signalling during recovery from severe injury

Our results show that the combined activities of regenerative states, FCC and revSCs, explain nearly all de novo crypt reconstitution in the irradiated intestines. Because chromatin remodelling also plays a crucial role in this process^22^, next we investigated the epigenetic landscape of the regenerating intestinal epithelium to understand how alterations in the epigenome enable reconstitution of fresh Lgr5^+^ CBC cells from revSCs. To do so, we performed single-nuclear multi-omics (i.e. single-nuclear RNA-seq and single-nuclear ATAC-seq) analysis of the recovering intestinal epithelium. We then compared these epitranscriptomes of irradiated intestinal epithelia from 0, 1 and 2 dpi with those from 4 dpi, a timepoint when fresh Lgr5+ CBC cells have just started to emerge^20,26^. We used weighted nearest neighbor (WNN) analysis, which integrates both gene expression and chromatin accessibility profiles. Cell types were identified using well-established gene cell markers including immune cells, fibroblasts, enteroendocrine (EE) cells, enterocytes (EC), goblet cells (GC), Paneth cells (PC), revival stem cells (revSCs), transit amplifying (TA) cells, and crypt base columnar (CBC) cells **(Fig. 6a, Extended Data Fig. 10a, and Supplementary Table 4)**.

**Fig. 6:**
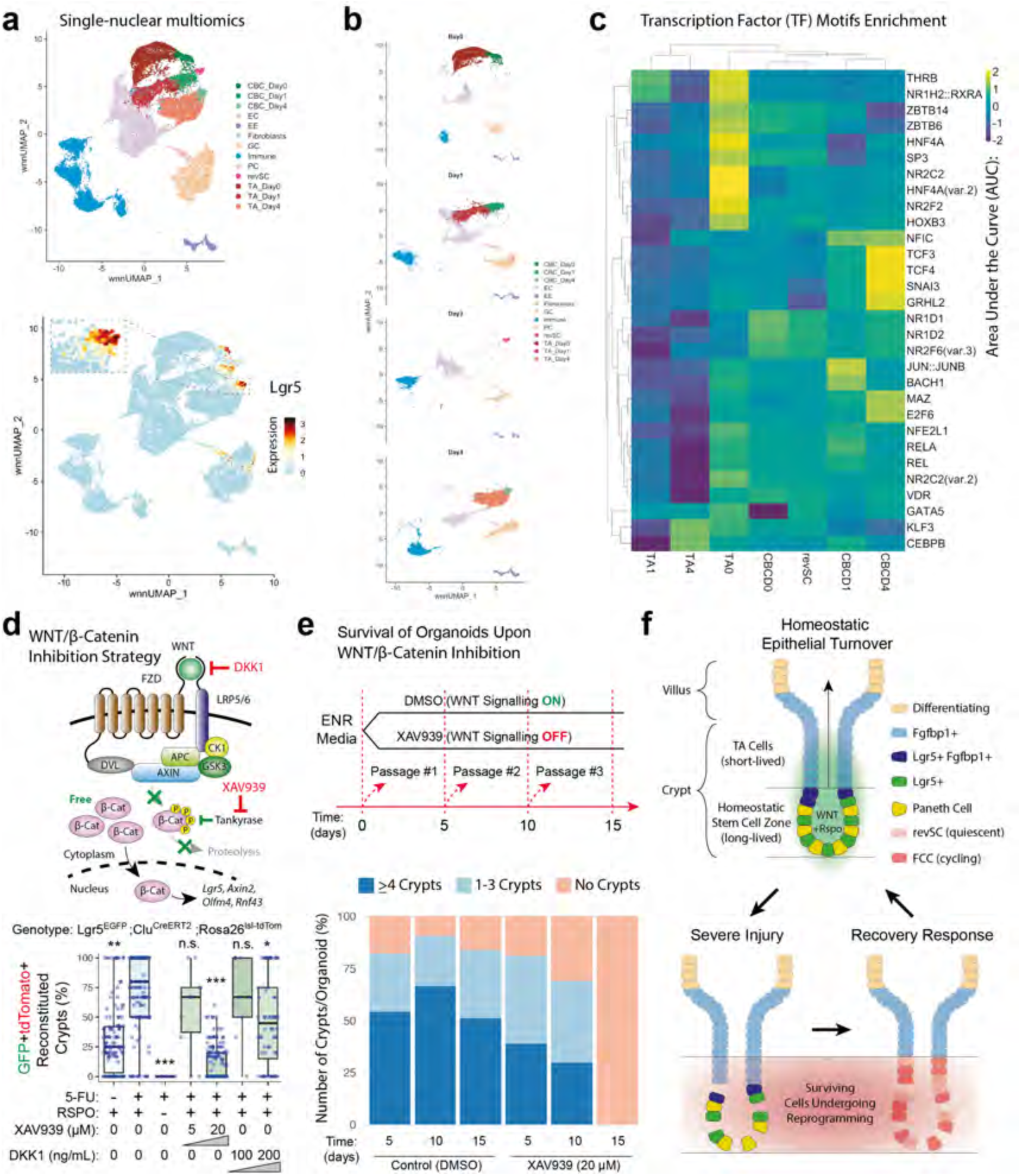
Integration of scRNA-seq and scATAC-seq reveals chromatin and signaling dynamics during intestinal regeneration and WNT pathway modulation **(a)** UMAP constructed containing scRNA and scATAC sequencing data using weighted nearest neighbor (WNN) analysis depicting intestinal cell type clusters before and after irradiation. **(b)** UMAP of WNN analysis at homeostasis (Day 0), 1 dpi, 2 dpi, and 4 dpi. **(c)** A heatmap of hierarchical clustering of highly expressed motifs for each cell cluster. **(d)** Schematic of WNT/b-catenin signalling when “ON” with depictions of incluence of DKK1 and XAV939 inhibitors (Top). The percentages of GFP+tdTomato+ reconstituted crypts from Lgr5DTR-EGFP and Clu^CreERT^^2^, Rosa26^loxp-STOP-loxp-tdTomato^ organoids with treatment combinations with 5-FU, RSPO, XAV939 and DKK1 and various concentrations of inhibitors (Bottom). **(e)** Organoid survival assay for 15 days with 3 passages and with or without XAV939 incubation. Number of crypts were counted from 0 to greater than 4 crypts depicted in shades of blue. **(f)** Schematic depicting homeostatic epithelial turnover (top) and regeneration response induced upon severe somatic injury (bottom).

Interestingly, the epigenetic profiles of both TA cells and Lgr5^+^ CBC cells changed dramatically following irradiation, resulting in time point–specific clustering of these populations **(Fig. 6a,b)**. By 2 dpi, nearly all CBCs and TA cells had been eliminated, and revSCs emerged as the predominant stem-like population **(Fig. 6b)**. Of note, FCCs were not detected in this dataset, potentially due to the low amount of mRNA transcripts that can be recovered from nuclei. By day 4, revSCs were no longer detectable, while both CBC cells and TA cells re-emerged to repopulate the epithelium. These findings align with previous reports identifying revSCs as a key epigenetic state that drives intestinal epithelial regeneration following irradiation through conversion to Lgr5^+^ CBCs^22^.

Next, we performed an unbiased differential motif enrichment analysis to identify transcription factor (TF) motifs with increased activity across epithelial cell types during regeneration **(Supplementary Table 5)**. Hierarchical clustering of motif area under the curve values revealed that Tcf3 and in particularTcf4 motifs displayed the highest accessibility in emerging CBCs at 4 dpi. Additional motifs, including Tcf3, Snai3, and Grhl2, were also enriched in CBCs at this time point, suggesting a potential role for these transcription factors in re-establishing epithelial homeostasis **(Fig. 6c** and **Extended Data Fig. 10b,c)**.

Interestingly, Tcf4 is a major target of β-catenin and has previously been shown to be essential for maintaining intestinal stem cell identity and driving proliferative activity in the intestinal epithelium^39^. These findings suggest that revSC to Lgr5^+^ CBC conversion may be driven by WNT/β-cateninsignalling. To test this notion, we switched to an intestinal organoid model that measures the role of fetal-like revSCs in reconstituting Lgr5^+^ CBCs following injury^22,26^. We used intestinal organoids that expressed *Lgr5-DTR-EGFP* and a TAM-inducible revSC-specific *Clu^CreERT^*^2^*; Rosa26^loxp-STOP-loxp-tdTomato^* reporters^26^. First, we incubated TAM-treated *Lgr5DTR-EGFP; Clu^CreERT^*^2^*, Rosa26^loxp-STOP-loxp-tdTomato^* reporter organoids with a DNA-damaging therapeutic agent 5-Fluorouracil (5-FU), to mimic radiation-induced tissue injury. This approach traced ∼80% of organoid crypts before physical dissociation **(Fig. 6d** and **Extended Data Fig. 10d)**. Treatment with WNT antagonists XAV939 and DKK1 significantly reduced the proportion of tdTomato^+^ de novo crypts in a dose-dependent manner, while no crypts were formed in the absence of R-spondin1. Importantly, neither XAV939 nor DKK1 treatments prevented early organoid growth (**Extended Data Fig. 10e**). In addition, IF imaging at 5 days after plating showed formation of de novo crypts that were not traced but contained abundant proliferating Lgr5^+^ and Ki67^+^ cells **(Extended Data Fig. 10d)**, indicating they did not arise from Clu^+^ revSCs.

However, organoids treated with the WNT antagonist XAV939 declined significantly after the second passage and could not be sustained beyond the third, indicating that WNT signalling is necessary not only for reconstituting the homeostatic stem-cell compartment during recovery from injury but also for maintaining basal intestinal epithelial turnover (**Fig. 6e** and **Extended Data Fig. 10e**).

In conclusion, our data show that epithelial turnover throughout the small intestinal epithelium is maintained by Lgr5⁺ CBCs and establish that reactivation of canonical WNT signalling is required for the epigenetic conversion of regenerative cells into bona fide Lgr5⁺ CBCs following severe somatic injury, thereby reinstating Lgr5⁺ CBCs at the apex of the intestinal epithelial cell hierarchy **(Fig. 6f)**.

## DISCUSSION

Intestinal epithelial turnover has traditionally been conceptualized as a hierarchically organized process sustained by Lgr5⁺ CBC cells^2^. However, recent studies have introduced alternative models in which differentiated and Lgr5⁻ cells can dedifferentiate and contribute to repair after injury^40^. More recently, several studies have described transient fetal-like regenerative states that arise upon damage and can reconstitute the Lgr5⁺ CBC pool^41^. These findings have led to the belief that the intestinal epithelium possesses remarkable plasticity, raising fundamental questions about the identity, exclusivity, and resilience of the intestinal stem cell compartment.

By integrating single-cell transcriptomics, chromatin accessibility profiling, lineage tracing using a non-disruptive *Lgr5-CreERT2; Rosa26-LSL-tdTomato mouse* model, and spatial gene expression analyses, we provide compelling evidence that Lgr5⁺ CBCs are the dominant and quantitatively essential source of epithelial renewal under homeostatic conditions. Within just seven days, nearly the entire crypt–villus axis—including all major epithelial lineages except Paneth cells—is replenished by Lgr5⁺ cells, reaffirming the classical model and addressing prior limitations stemming from mosaic and hypomorphic reporter systems^2^.

Beyond confirming the centrality of Lgr5⁺ CBC cells, we have uncovered previously unappreciated heterogeneity within this population. We identify two transcriptionally and functionally distinct CBC subsets: CBC1, a relatively quiescent population localized at the crypt base with lower β-catenin activity, and CBC2, a more proliferative subset positioned slightly higher and enriched for WNT target gene expression. This internal stratification underscores the complexity of the Lgr5⁺ stem cell pool and may reflect differential responsiveness to niche-derived cues.

Recent proposals of Fgfbp1⁺ IPCs as a hierarchically upstream population that generates Lgr5⁺ cells^18,19^ are not supported by our data. We demonstrate that *Fgfbp1* expression is broadly distributed across TA and early progenitor cells, and while some Lgr5⁺ cells co-express *Fgfbp1*, the majority of Fgfbp1⁺ cells lack *Lgr5* expression and reside outside the stem cell zone. In addition, nearly 100% Fgfbp1^+^ cells are directly replenished by Lgr5^+^ cells within seven days.

These findings argue against a model in which Fgfbp1⁺ IPCs function as a distinct, upstream stem cell population and instead position them as downstream or lateral derivatives of canonical Lgr5^+^ CBC cells.

Following injury, when Lgr5⁺ CBCs are quantitatively depleted, we observe the rapid emergence of two transcriptionally diverging regenerative states: quiescent revSCs and proliferative FCCs. While both revSCs and FCCs express fetal-like genes such as Ly6a/Sca1, revSCs enter a quiescent state marked by *Clu* and a complete absence of CBC or cell cycle genes, whereas FCCs express *Lgr5*, *Ascl2*, *Fgfbp1*, and cell-cycle genes (e.g., *Mki67*) at low levels. In addition, revSC induction peaks at 3 dpi, and FCCs peak at 4 dpi. Trajectory analysis focusing on transitional states at 4 dpi posits that revSCs give rise to FCCs, which then reconstitute the CBC compartment. Consistent with this model, YAP signalling is progressively inhibited, whereas WNT/β-catenin signalling and proliferation are induced as cells transition from revSCs to FCCs and ultimately to homeostatic CBCs.

Despite the transient prominence of these regenerative populations, lineage tracing across early (7 days) and late (31 days) timepoints before irradiation demonstrates that Lgr5⁺ cells and their descendants ultimately re-establish the crypt architecture and sustain intestinal regeneration following different severities of radiation injury. This regenerative trajectory is critically dependent on reactivation of canonical WNT/β-catenin signalling. In both homeostasis and injury contexts, we find that WNT activity, for example, indicated by TCF motif accessibility and β-catenin target gene expression, is tightly associated with proliferative potential and stem cell function. Withdrawal of R-spondin or pharmacological inhibition of WNT signalling using XAV939 or DKK1 abolishes crypt formation and severely impairs regenerative output.

Together, our findings use quantitative approaches to reaffirm the position of Lgr5⁺ stem cells at the top of the cell hierarchy throughout the small intestinal epithelium and redefine epithelial repair following somatic injury as a biphasic process: an initial WNT-independent induction of fetal-like states, followed by a WNT-dependent reconstitution of the Lgr5⁺ stem cell compartment. This framework reconciles previously conflicting models by demonstrating that facultative plasticity and fetal-like reprogramming are not alternatives to Lgr5⁺ stem cell function, but transient compensatory states that ultimately restore Lgr5⁺ identity and the canonical WNT-driven regenerative program^41^. In doing so, we establish Lgr5⁺ CBCs and WNT/β-catenin signaling as indispensable drivers of both intestinal homeostasis and repair.

## Resource availability

### Correspondence

Further information and requests for resources and reagents should be directed to the corresponding authors.

### Materials availability

Any material that can be shared will be released via a material transfer agreement for non-commercial usage.

### Data and code availability

- All data from this study are available from the corresponding authors upon reasonable request. Single-cell RNA and ATAC-seq generated from this study can be found at GEO under accession ID GSE304776.
- Codes for original analysis can be found here: https://github.com/jchua2468/Chua_AyyazLab_2025_Wntable
- Any additional information required to reanalyse the data reported in this paper is available from the lead contact upon request.

## METHODS

### Animal models

All procedures with mice were approved by the Institutional Animal Care and Use Committee (IACUC) at Duke University and the Canadian Council on Animal Care. *Lgr5-CreER* mice were generated by Huch and colleagues^25^, and *Rosa26-LSL-tdTomato* (Ai9) mice^42^ were purchased from the Jackson Laboratory (Strain #:007909). Mice were at least 8 weeks old at the time of irradiation. Both sexes of mice were used, and littermates were used as controls for all experiments.

### Intestinal Organoid Culture

Small intestinal organoid cultures were established following a previously described method^22^. Briefly, the mouse small intestine was incubated in PBS with 2mM EDTA and epithelial fragments were filtered through a 70 μm cell strainer to isolate crypt cells. These crypt cells were cultured in 35 μl of Matrigel (BD Biosciences) and grown in media containing Advanced-DMEM/F12 (Life Technologies), B27 supplement (Life Technologies), 2mM Glutamax (Life Technologies), 100 U/mL Penicillin/100 mg/mL Streptomycin (Life Technologies), N2 supplement (Life Technologies), mouse recombinant Egf (Life Technologies), 100 ng/mL mouse recombinant Noggin (Peprotech), and 300 ng/mL human R-spondin1 (R&D Systems).

### Immunofluorescence (IF)

Mouse small intestines were harvested, washed with ice-cold PBS, and cut into smaller fragments. These pieces were fixed overnight in 10% buffered formalin, followed by overnight transfer to 30% sucrose PBS at 4°C for cryoprotection. The tissue fragments were then stored in OCT at -80°C for long-term preservation. OCT blocks were sectioned into 16 μm thick slices, permeabilized with 0.5% Triton X-100 and 0.2% Tween 20 in PBS for 30 minutes and blocked in 2.5% BSA for 1 hour. Sections were incubated overnight with primary antibodies targeting the following markers: mouse monoclonal anti-Ki67 (BD Biosciences, Cat# 55600), goat polyclonal anti-GFP (Abcam, Cat# ab5450), rabbit monoclonal anti-Olfm4 (Cell Signaling Technology, Cat# 39141S), goat anti-Sca1Ly6 (R&D Systems, Cat# AF1226-SP), goat anti-Lrig1 (R&D Systems, Cat# AF3688), rabbit anti-Fzd5 (Abcam, Cat# ab75234) and rabbit polyclonal anti-lysozyme (Dako, Cat# EC32117). After primary antibody incubation, secondary antibodies conjugated with DyLight fluorophores (Fisher Scientific) were applied, and sections were counterstained with DAPI (Sigma-Aldrich) for 30 minutes. Images were acquired at 10X magnification using a Leica fluorescent microscope and at 63X magnification on a Nikon Ti2 confocal microscope.

### Regeneration Experiments

#### In vivo

Mice were exposed to a single dose of sub-total-body irradiation (SBI) with the head and forelimbs shielded to prevent hematopoietic acute radiation syndrome^37^. Jigs were used to hold unanesthetized mice to limit movement during radiation exposure. SBI was performed using the CIX3 small animal irradiator (Xstrahl) at Duke University. Mice were placed on a turntable shelf and treated 40 cm from the radiation source. A total physical dose of 10, 12, or 14 Gy X-rays was delivered using 320 kV, 10 mA, and 1mm Cu filter. The dose rate was 3.44 cGy/ sec. The radiation dosimetry was performed using Thermo Luminescent Dosimeters (TLDs) placed inside tissue equivalent mouse phantoms. The TLDs were cross calibrated using an ion chamber that had been calibrated with a NIST traceable X-ray calibration source.

### Tissue Histology

Irradiated and unirradiated mice were sacrificed by CO_2_ asphyxiation at indicated time points, where the proximal 12 cm of the jejunal region of the small intestine was isolated and flushed with cold DPBS (Gibco). To generate formalin-fixed paraffin-embedded blocks, tissues were divided into 2 cm segments and fixed in 10% neutral buffered formalin for 24 hours, then transferred to 70% ethanol for up to one week prior to processing.

### RNA in situ hybridization (RNAScope)

RNAscope in situ hybridization using RNAscope Multiplex Fluorescent V2 detection kit (ACD, 323110) was performed according to manufacturer’s protocol with modifications. Paraffin sections (10 μm thickness) were deparaffinized, then incubated in 3% H_2_O_2_ in PBS with 0.1% Tween 20 (PBS-T) for 30 min at RT. After rinsing slides with PBS-T briefly, sections were heat-decrosslinked in 10 mM Tris, 1 mM EDTA (pH 9.0) around 95 °C for 15 min. Slides were cooled down by dipping in PBS-T at RT, then incubated in pre-warmed PBS-T containing 0.2 μg/ml proteinase K (NEB, P8107S) at 37 °C for 30 min. After washing slides with PBS-T 3 times, slides were incubated with RNAscope probes according to manufacturer’s protocol. We used probes at a lower concentration than to manufacturer’s suggestion, to minimize signal crosstalk. Catalog numbers of probes and actual dilutions used in this study are shown in List of Reagents. Washes after incubation with probes, AMP1, AMP2, AMP3, HRP-C1, HRP-C2, HRP-C3 or HRP-C4 were carried out using 0.5x SSC, 0.1 % Tween 20 for 5min x2 at 37C. Other washes were done with PBS-T at RT. For signal development, TSA Vivid Fluorophore Kit 520, 570, and 650 from Bio-Techne (Cat. No. 7523, 7526 and 7527) and iFluor 750 styramide from AAT Bioquest (Cat. No. 45065) were used. DAPI (Simga-Aldrich, D9542) was used for nuclear counterstain at 0.5 μg/ml in PBS-T.

Images were acquired using either a Zeiss Axio Observer Z1 inverted microscope with a modified Yokogawa CSU-X1 spinning disc confocal scanhead, or a Nikon Ti2 inverted microscope with AXR confocal scanner. Nikon NIS-Elements software was used for image processing.

### Quantification of lineage tracing images

*Lgr5-CreERT2; Rosa26-LSL-tdTomato* mice received either sham IR, 10, 12, or 14 Gy SBI. To induce Cre-ER-mediated recombination, mice were given two injections of 100mg/kg Tamoxifen (Sigma) intraperitoneally at various time points either prior to sham IR or SBI. Surviving mice were euthanized 10 days post-SBI or sham-IR for tissue harvest. Small intestines (duodenum, jejunum, ileum segments) were harvested and fixed in 4% paraformaldehyde (PFA) in 1X PBS for 2 hours, then transferred to 30% sucrose in PBS overnight at 4C. After fixation, intestines were swiss rolled luminal side up from proximal to distal end and frozen in OCT^43^. Samples frozen in OCT were cut to 10 μm sections and mounted onto charged slides. Tissue slides were fixed with 4% PFA in 1X PBS for 10 minutes, briefly washed with 1X PBS, and mounted with ProLong™ Diamond Antifade Mountant with DAPI (Invitrogen). Images were taken at 10X for crypt quantification, 20X for representative crypt images, and 2.5X for representative full Swiss-rolled images using a Leica fluorescent microscope.

A total of 10 representative 10X images from proximal duodenum to distal ileum were taken for quantification purposes for each slide. For experiments harvested at time of sham-irradiation (harvested at day 0), tdTomato^+^ cells within intestines were separated into 5 regions from crypt to villus for quantification purposes. Region 1 was denoted as tdTomato^+^ cell position 0 to +4, region 2 was denoted as tdTomato^+^ cell position +5 to +8, region 3 was denoted as tdTomato^+^ cell position +9 to below crypt-villus junction (CVJ), region 4 was denoted as tdTomato^+^ cell position at CVJ, and region 5 was denoted as tdTomato^+^ cell positions above the CVJ, such as in the villus. 100 crypts were quantified per mouse and averaged per region. For 10-day post-IR experiments, crypts were quantified by counting the number of crypts that have ≥ 50% tdTomato^+^ signal. The percentage of lineage tracing events per slide was determined by dividing the total number of positive tdTomato crypts by the total number of crypts present.

### scRNA Reference Pre-processing

Previously published single-cell sequencing files^6,20,22,24^ including GSE files GSE123516, GSE230766, GSE92332, GSE201859, and this study (GSE304776) were extracted and converted into Seurat objects using Seurat version 4.0.1 and corresponding metadata was added^44^. Samples were filtered to have cells containing over 200 transcripts, less than 6000-8000 transcripts, and less than 10-40% mitochondrial DNA. To account for sample-specific variability in sequencing depth and quality, filtering thresholds were adjusted per sample rather than uniform adjustments. Thresholds were tailored based on sample mean and distribution. All samples were then merged into one main object and the top 2000 variables genes, and 50 principles component were used to run principal component analysis (PCA). To address batch effect among samples from various sources we employed the Harmony package for our analysis^45^. Harmony integration was performed using default parameters to merge five embeddings. Principle ss 1 through 35 were used for clustering visualization, with Louvain clustering applied at a resolution of 0.1.

Cell type markers previously used in^20^ were used to identify cell types corresponding to each cluster. Immune related clusters were removed from the dataset to leave only epithelial cells.

### Cell type Identification and distribution during regeneration

Cell type makers for the intestine previously used in^20^ were plotted on the scRNA data to look at average log2 expression. These values were converted to a dot plot to visualize overall expression in a cell type, as well as percentage of cell expression.

Cell types were identified and converted to percentages based on their proportion of the whole scRNA sample. This allowed analysis of cell contribution of the whole intestinal sample. These distribution patterns were also calculated and visualized in pie charts to see how cell contribution dynamics change over the course of intestinal repair and regeneration. This was done for both the reference and query dataset. These changes in cell contribution over time were then compared between each other to calculate fold change of cell types for Day 1, Day 2, Day 3, and Day 4 for the reference dataset.

### scRNA Query Clustering

Combined scRNA datasets were extracted and converted into a Seurat object as described earlier. Samples were filtered to contain cells with >200 transcripts, <7500 transcripts, and <25% mitochondrial DNA. To follow parallel with the reference dataset 2000 variable genes, and 50 principle components were used to run PCA analysis. The Harmony package was also used with default parameters to merge five embeddings. Principle components 1 through 35 were used for clustering visualization, with Louvain clustering applied at a resolution of 0.5.

For running query visualization on the reference dataset, core anchors in the reference dataset were identified for each cell type based on 1 to 30 dimensions. Predictive anchors based on reference anchors were identified in the query dataset and assigned to each cell. The query dataset was then visualized based on reference dataset cell type.

### scMULTIOMICS Sample prep

Nuclei were extracted by resuspending crypt samples in 100 µL of chilled lysis buffer (10X Genomics Protocol CG000365 Rev C) for 5 minutes on ice, washed twice with lysis buffer, resuspended at 5,000 nuclei per µL and processed with the 10X genomics Multiome ATAC + Gene Expression kit targeting 10,000 nuclei per sample and sequenced on NovaSeq6000.

### scMULTIOMICS Pre-processing using Weighted Nearest Neighbor

To process fastq files for ATAC and RNA sc data mm39 reference genome was used with CellRanger ARC 2.0 (10X genomics) with default settings. RNA outputs from cell ranger were converted into Seurat objects and ATAC outputs were constructed into a global peak set using corresponding fragment files of the reference genome. ATAC sequencing was then ranges were then annotated using ensemble mouse database version 79 to map peaks to gene locations.

Both scRNA assays and scATAC assays were merged for each sample separately. All datasets were merged and filtered to remove samples that had <700,000 ATAC fragments, >1000 ATAC fragments, >25,000 RNA transcripts, > 500 RNA transcripts, and < 40% mitochondrial DNA. For the scRNA assay we normalized and identified variable features using SCTransform and used dimensions 1 to 50 for independent visualization. For the scATAC assay we normalized the data using TF-IDF and SVD, which is tailored for sparse high-dimensional data like ATAC sequencing. For independent visualization dimensions 2:50 were used. The first dimension was excluded as it commonly captures technical variance in scATAC sequencing data.

Seurat’s Weighted Nearest Neighbor was utilized to merge and analyze scRNA and scATAC sequencing data^44^. Multimodal Neighbors were identified between scRNA and scATAC sequencing using PCA and 1 to 25 dimensions for the scRNA assay and LSI and 2 to 25 dimensions for the scATAC assay. A resolution using the Louvain clustering of 0.25 was used. Gene markers as stated above were used to identify cell identities to each cluster.

### Crypt sub clustering and merging

Clusters corresponding to cells located in the crypt region of the intestine were identified and subclustered separately from the original object to isolate CBCs. scRNA and scATAC assays were reanalyzed using the same parameters as stated earlier, but a resolution of 0.9 was used. CBCs and other crypt related cell types were identified using markers stated previously. The new cell type identity in the sub clustered data was added to the metadata of the original object based on barcode identity. The original object with these annotations was then visualized for downstream analysis.

### Differential Motif activity interference and integration with gene expression

To quantify transcription factor motif activity across single cells, we performed motif accessibility analysis using Signac and chromVAR^46,47^. Position weight matrices for human transcription factors were retrieved from the JASPAR2020 database species 9606. Although the scATAC sequencing of interest was from mouse we used human position weight matrices to enable broader mapping of evolutionary conserved transcription factor motifs. Motif annotations were added by creating a motif matrix correlating with peaks in the scATAC data and the mm10 genome. The resulting data was stored within the ATAC assay within the multiomics data. To estimate transcription factor motif accessibility per cell we ran chromVAR using the BSgenome.Mmusculus. UCSC.mm10 genome.To identify possible transcription factors corresponding to each cell type we applied a Wilcoxon rank-sum test implemented in the presto package^48^. Differentially accessible motifs were computed from chromVAR scores and differentially expressed genes were computed from SCTransform-normalized RNA data. For both modalities candidates from each cell type were screened based on log fold change, AUC, and adjusted values. Motif IDs were also converted to gene names to remove multiple isoform suffixes.

### Identification of cell type cell cycle phase distribution

Cell cycle marker genes were obtained through a previous publication that marked cells in G1, G2M, and S phase^49^. These markers were plotted on the reference and query scRNA data and cells were identified to be one of the cell cycle phases based on their highest log2 average expression cell cycle profiles. For each individual cell type, the number of cells in each phase of the cell cycle was converted to a percentage of cells contributing to the specific cell type. The percentage values of each cell type were then used to compare cell cycle dynamics within each cell type.

### WNT signalling cell distribution and cell expression markers

For each cell type various markers such as Lgr5, Fgfbp1, and cycling signatures were plotted as violin plots for expression level and cell distribution. Positive Lgr5 high and Fgfbp1 high cells (set threshold of) were also calculated using absolute values to display positive cell correlations for each cell type.

Znrf3, Rnf43, and β-catenin signals identified by a previous paper were plotted in a dot plot for each cell type. This visualization was to indicate what cell types had elevated levels of Wnt signalling, and what portion of cells were positive.

### Identification of gene markers for FCCs and revSCs

Sca1 were ploted and cells with an average log2 value of over 1 were Sca1 positive. These cells were then classified into FCCs and revSCs based on proliferative markers and cycling markers. Various genes were ploted between FCCs and revSCs and a paired T test was used to compare differences in FCCs and revSCs that contained these markers.

### Normalized expression

Reference and query datasets were normalized independently to account for read depth variety. Various genes transcript quantities were then calculated for each dataset at different timepoints to identify differences in specific gene transcript levels overtime.

### Velocity trajectory analysis

RNA velocity analysis was performed on the mouse intestinal epithelial using the scVelo (v0.2.5) pipeline^27^. Loom-formatted spliced/unspliced counts were loaded and filtered with a minimum shared count threshold of 20 and 2,000 highly variable genes. PCA neighborhood calculation and UMAP embeddings were computed using Scanpy and 30 PC and neighbors were used^50^. Velocities were inferred using stochastic modelling and UMAP embeddings from previous reference map were used as a projection.

### Generating Schematics

Publishing licensure was obtained for all images created with BioRender by L.D. through a Duke University premium subscription. Publication agreement numbers can be found below:

**Figure 2B-D:** Publication and Licensing Rights Agreement # DM28MJ3O1G

**Figure 2E:** Publication and Licensing Rights Agreement # LW28MJ345L

**Figure 5A:** Publication and Licensing Rights Agreement # WU28MJ3XFP

**Figure 5B:** Publication and Licensing Rights Agreement # AS28MJ41JI

**Figure 5E:** Publication and Licensing Rights Agreement # QO28MJ45G1

## Acknowledgements

We thank Dr. Hans Clevers for providing *Lgr5-CreER* mice. This study was supported by the following research funds: National Health Institute (USA) grants U01AI186969 (D.G.K, J.R. and C-L.L), U19AI067798 (D.G.K. and C-L.L), R21AI193496 (C-L.L), and Natural Sciences and Engineering Research Council of Canada (NSERC) Discovery Grant RGPIN-2024-04264 (A.A.). Whitehead Scholar Award from Duke University School of Medicine (C-L.L.) and Canadian Institutes for Health Research (CIHR) Doctoral Research Award (J.C.).

## Authors contributions

J.C., L.D., C.L.L., J.L.W., and A.A. co-conceived the study. J.C., L.D., M.N., M.F., L.L., R.H., and J.T. performed experiments; J.C., A.K. and A.A. conducted bioinformatic analyses; J.R., D.G.K., C-L.L., J.L.W., and A.A. acquired funding; J.C., L.D., C-L.L., and A.A. wrote the original draft; All authors reviewed and edited the final manuscript; C-L.L. and A.A. contributed equally to this manuscript.

## Supplementary Figures

**Extended Data Fig. 1:**
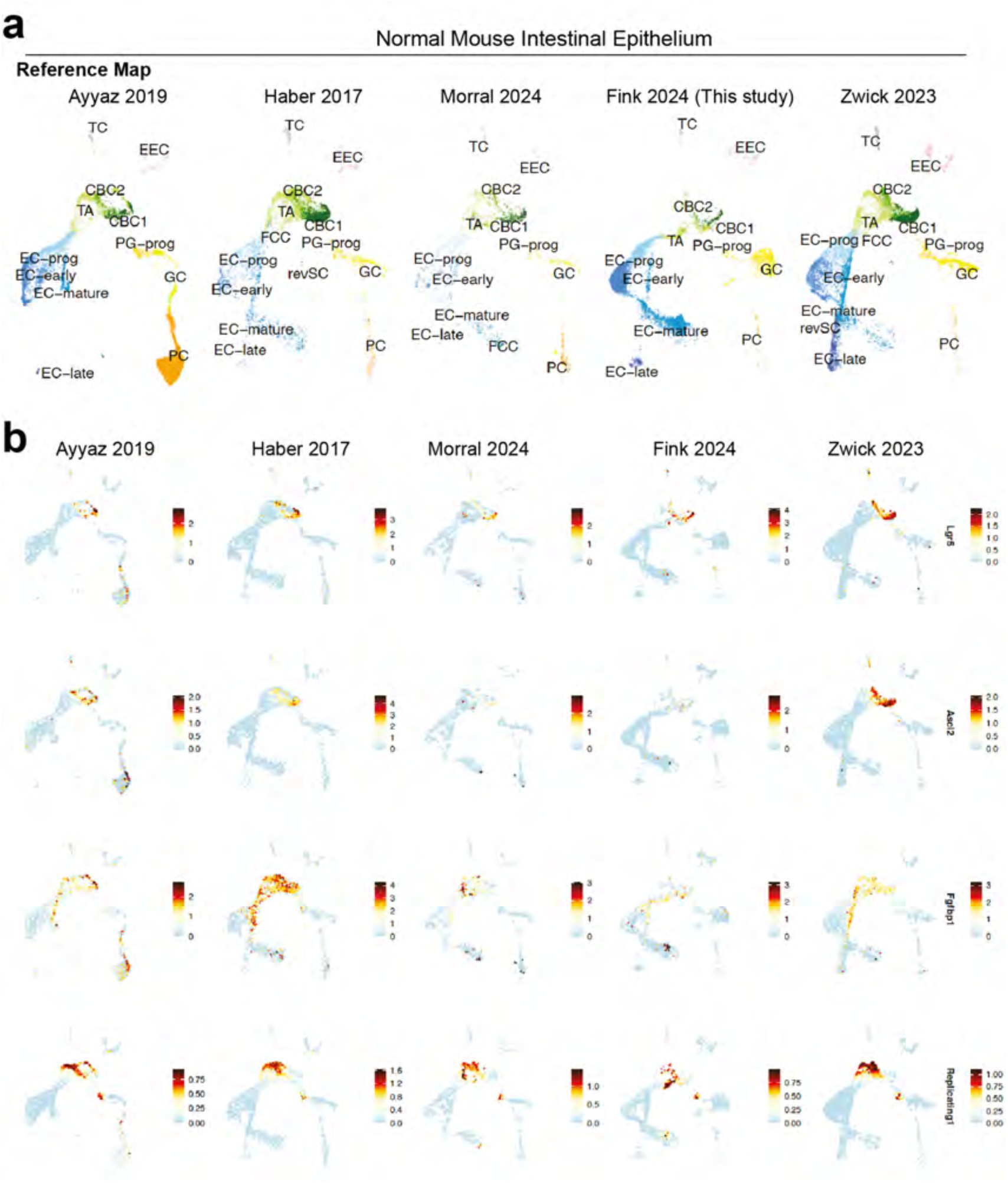
Identification of cell types across homeostatic murine intestine datasets **(a)** Separate UMAPs for 5 different datasets that comprise the reference dataset or a normal mouse intestine. **(b)** Feature plots for all 5 datasets for Lgr5, Ascl2, Fgfbp1, and replicating cell markers. Legend indicates average Log2 expression.

**Extended Data Fig. 2:**
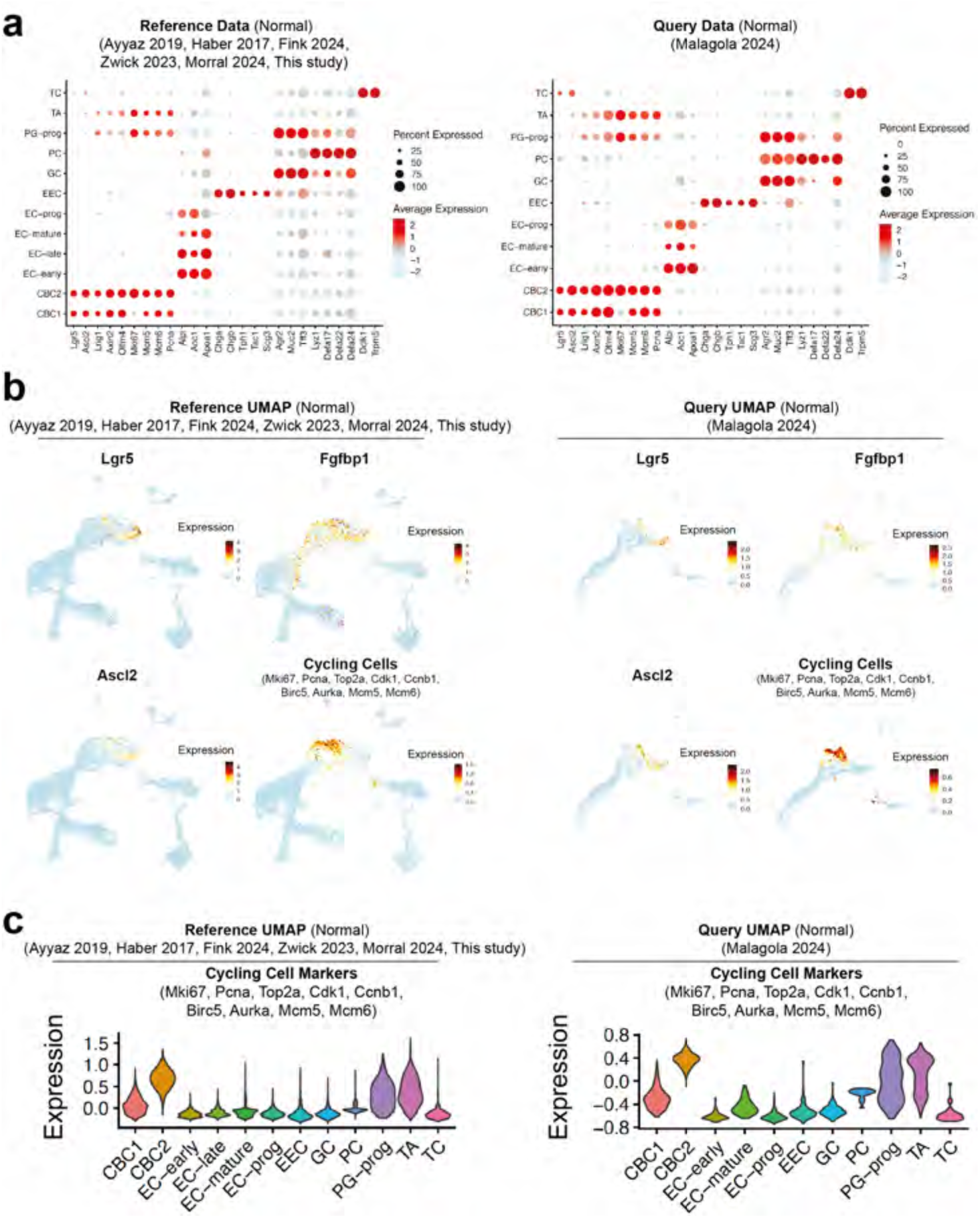
Cell type comparison and characterization between reference and query datasets **(a)** A dot plot depicting percentage of expressing cells and average expression for various intestinal stem cell markers. Left, indicates values for the reference dataset, and right, indicates values for the query dataset. **(b)** Feature plots for Lgr5, Fgfbp1, Ascl2, and cell cycling genes for left, reference dataset, and right, query dataset. Legend indicates average Log2 expression. **(c)** Violin plots showing expressing pattern of indicated cell cycle gene signature across two datasets.

**Extended Data Fig. 3:**
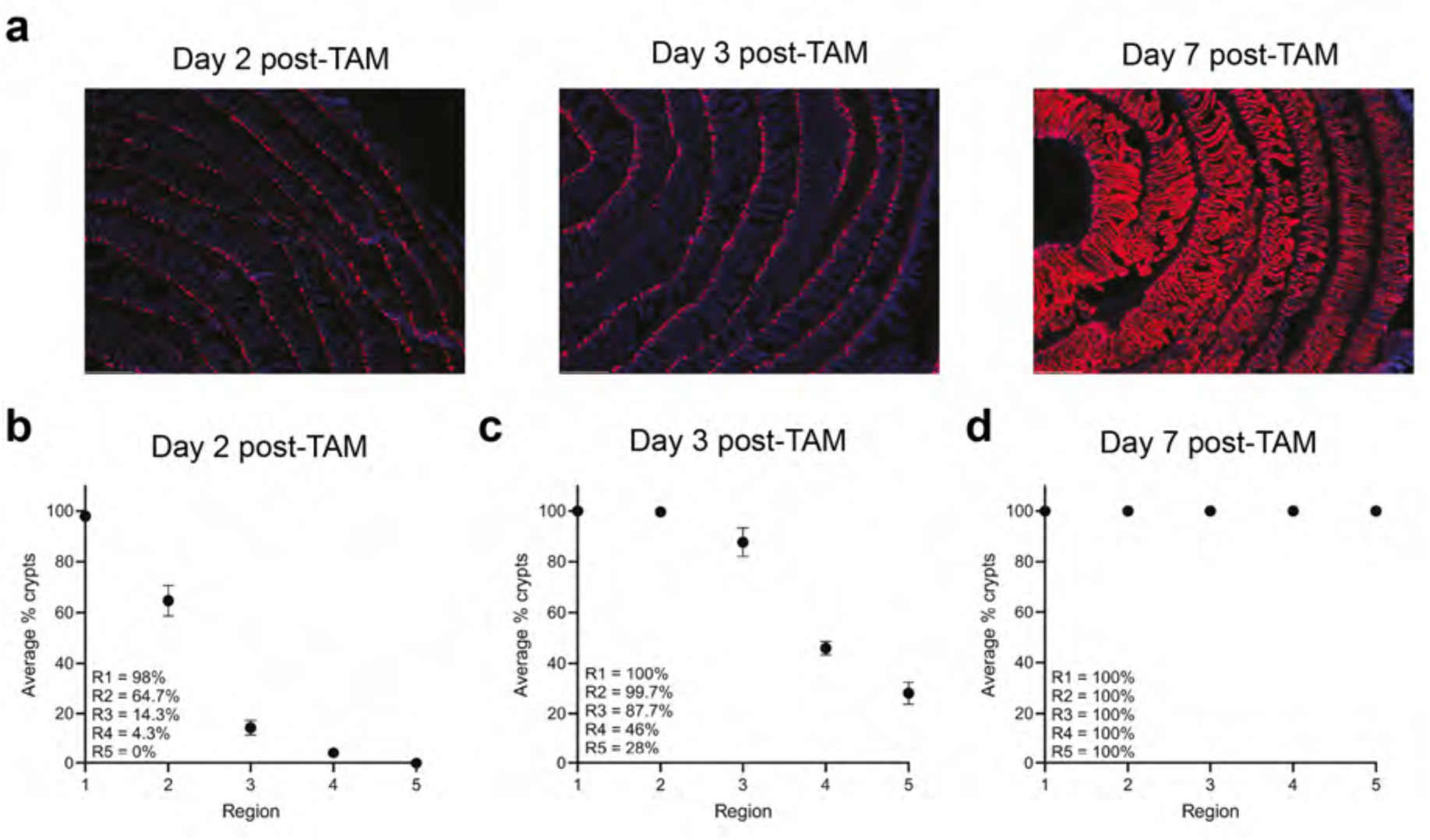
Lineage tracing reveals dynamics of Lgr5+ cells in unirradiated intestine epithelium **(a)** Representative 2.5X images of unirradiated Lgr5-CreERT2; Rosa26-LSL-tdTomato small intestines when given TAM to label Lgr5 progeny for 2 days, 3 days or 7 days prior to harvest. Scale bar 500um. **(b-d)** Quantification of 2 day, 3 day, and 7 day TAM labeling unirradiated Lgr5 cell positions/region. Each dot represents an average of each region from n=3 mice and n=300 total crypts quantified/graph.

**Extended Data Fig. 4:**
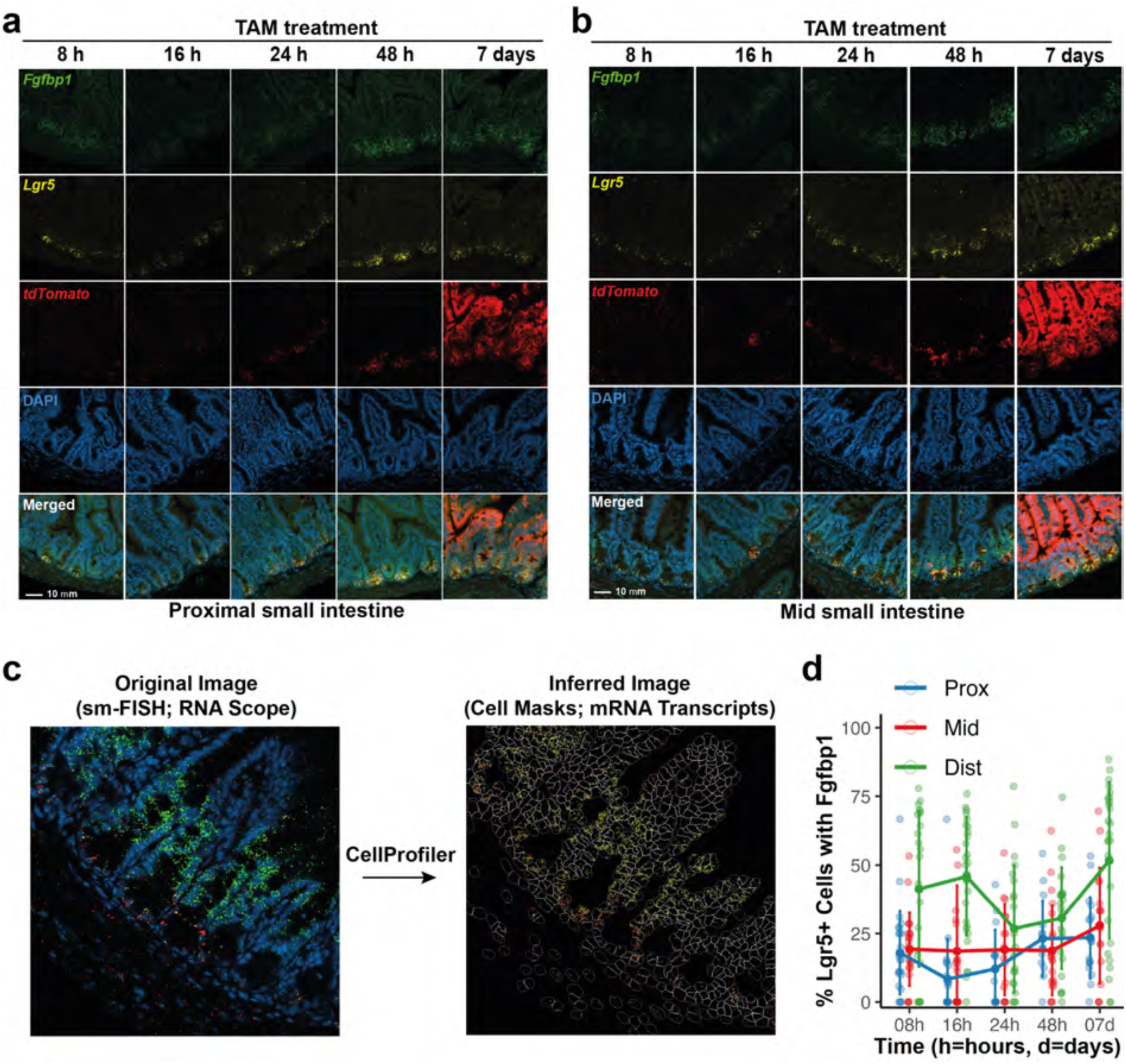
Early dynamics of tdTomato labeling in the small intestine and spatial analysis of Lgr5 and Fgfbp1 **((a)** Detection of tdTomato transcripts in the mid small intestine at an early phase after TAM administration. RNAscope in situ hybridization with Fgfbp1, Lgr5 and tdTomato probes on proximal small intestine sections treated with TAM from 8 h to 7 days. Region 1 (from 0 to+4) is indicated with dotted lines. **(b)** Detection of tdTomato transcripts in the proximal small intestine at an early phase after TAM administration. RNAscope in situ hybridization with Fgfbp1, Lgr5 and tdTomato probes on proximal small intestine sections treated with TAM from 8 h to 7 days. Region 1 (from 0 to+4) is indicated with dotted lines. **(c)** RNAscope images were processed through a CellProfiler pipeline to generate cell masks and assign transcripts to each cell. **(d)** A line plot showing the percentage of Lgr5⁺ cells expressing Fgfbp1 across proximal, mid, and distal regions over time.

**Extended Data Fig. 5:**
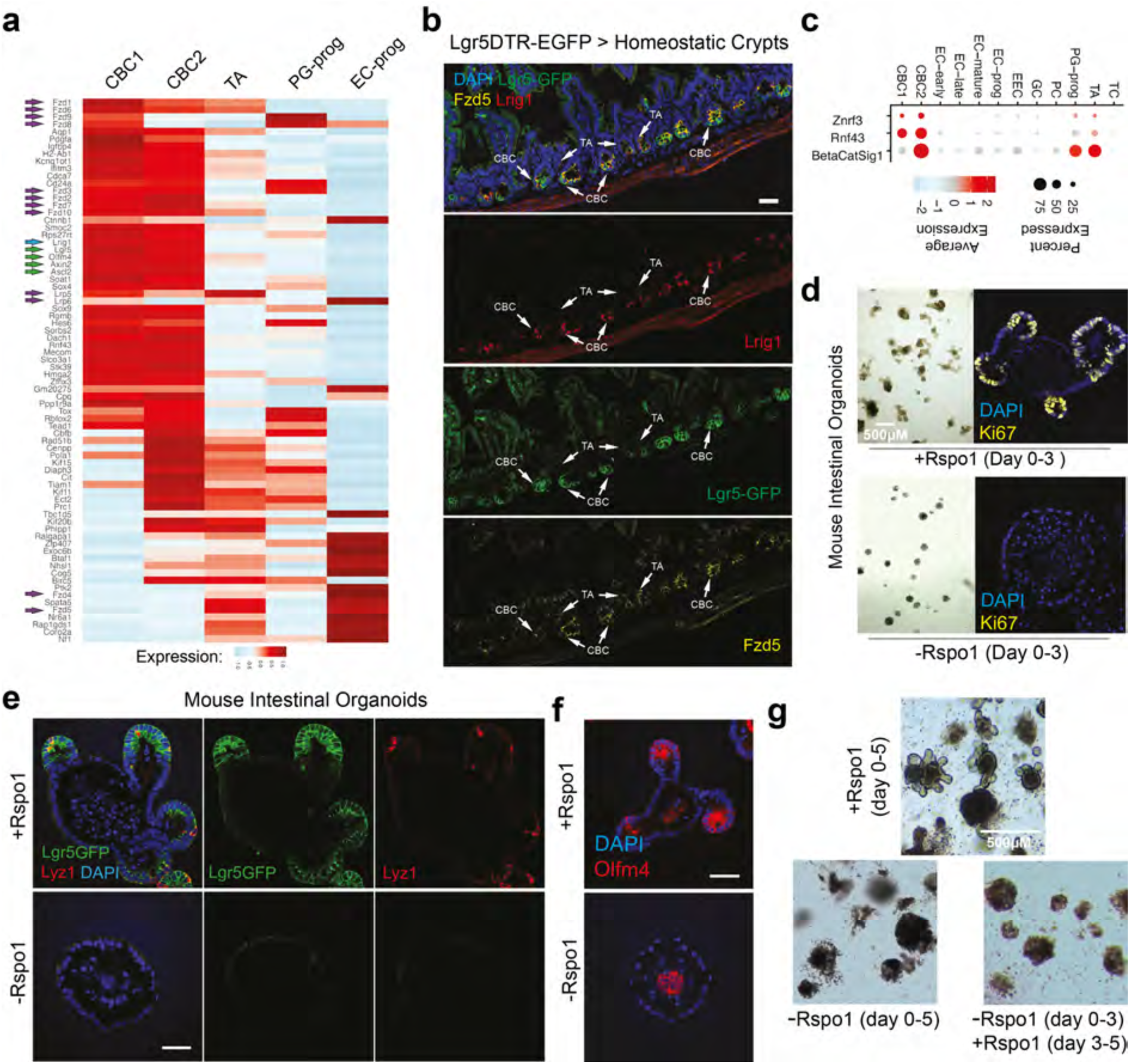
Spatial analysis of proliferation and WNT signaling in the small intestine **(a)** Heatmap displaying expression of stemness-associated genes in CBC1, CBC, TA, Paneth/enteroendocrine progenitors (PG-prog), and enterocyte progenitors (EC-prog) Purple arrows highlight Wnt-related genes, green arrows denote CBC-associated genes, and blue arrows indicate quiescent cell associated genes. **(b)** Immunofluorescence of Lgr5DTR-EGFP mouse intestines stained with Lrig1 and Fzd5. Arrows (white) indicate CBC and TA cell compartments. **(c)** Dot plot summarizing both expression level and percentage of cells expressing beta-catenin signaling pathway genes across all epithelial cell types. **(d)** Brightfield and immunofluorescence images of mouse intestinal organoids with and without the presence of Rspo1. Organoids stained with DAPI and Ki67. **(e)** Confocal images of Lgr5-GFP intestinal organoids, with and without Rspo1, stained for DAPI and Paneth cell marker Lyz1. **(f)** c57 mouse derived intestinal organoids cultured with (left) and without Rspo1 (right). Organoids were stained with DAPI and Olfm4. **(g)** Representative images of organoids cultured either continuously in Rspo1-supplemented or Rspo1-deficient medium for 5 days, or subjected to 3 days without Rspo1 followed by 2 days with Rspo1.

**Extended Data Fig. 6:**
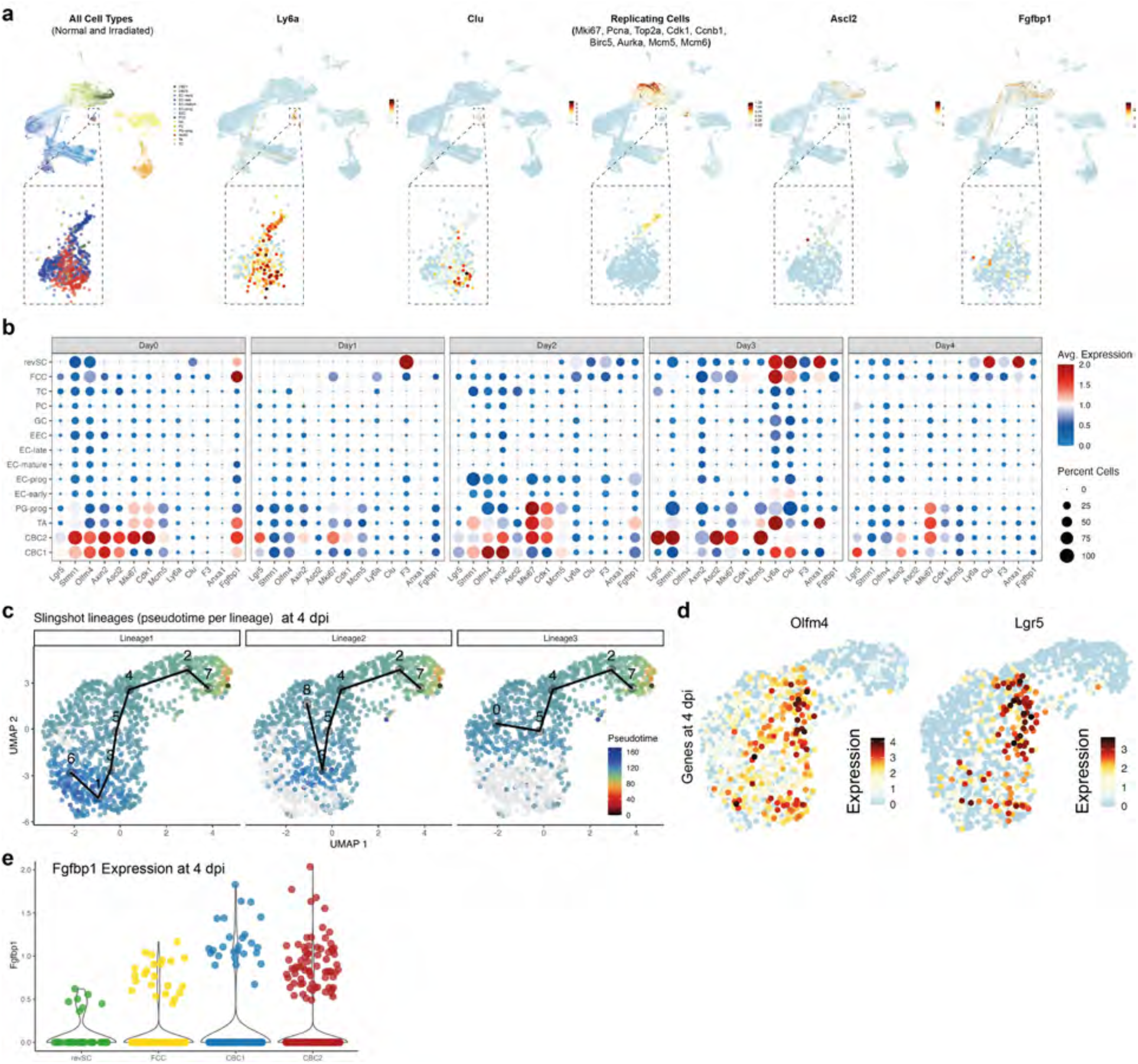
Gene expression dynamics of fetal-like program **(a)** UMAP plots showing indicated gene expression patterns across all time points. **(b)** Dotplot showing alterations in homeostatic stem and regenerative cell-specific gene expression patterns across indicated time points. **(c)** Three lineages revealed by Slingshot based pseudotime analysis at 4 dpi are separately plotted on UMAP plots. **(d)** Expression of CBC markers Olfm4 and Lgr5 is plotted on UMAP using single-cell transcriptomes of intestinal epithelium recovering from irradiation at 4dpi. **(e)** Fgfbp1 expression is plotted on a violin plot across stem cell compartment at 4 dpi.

**Extended Data Fig. 7:**
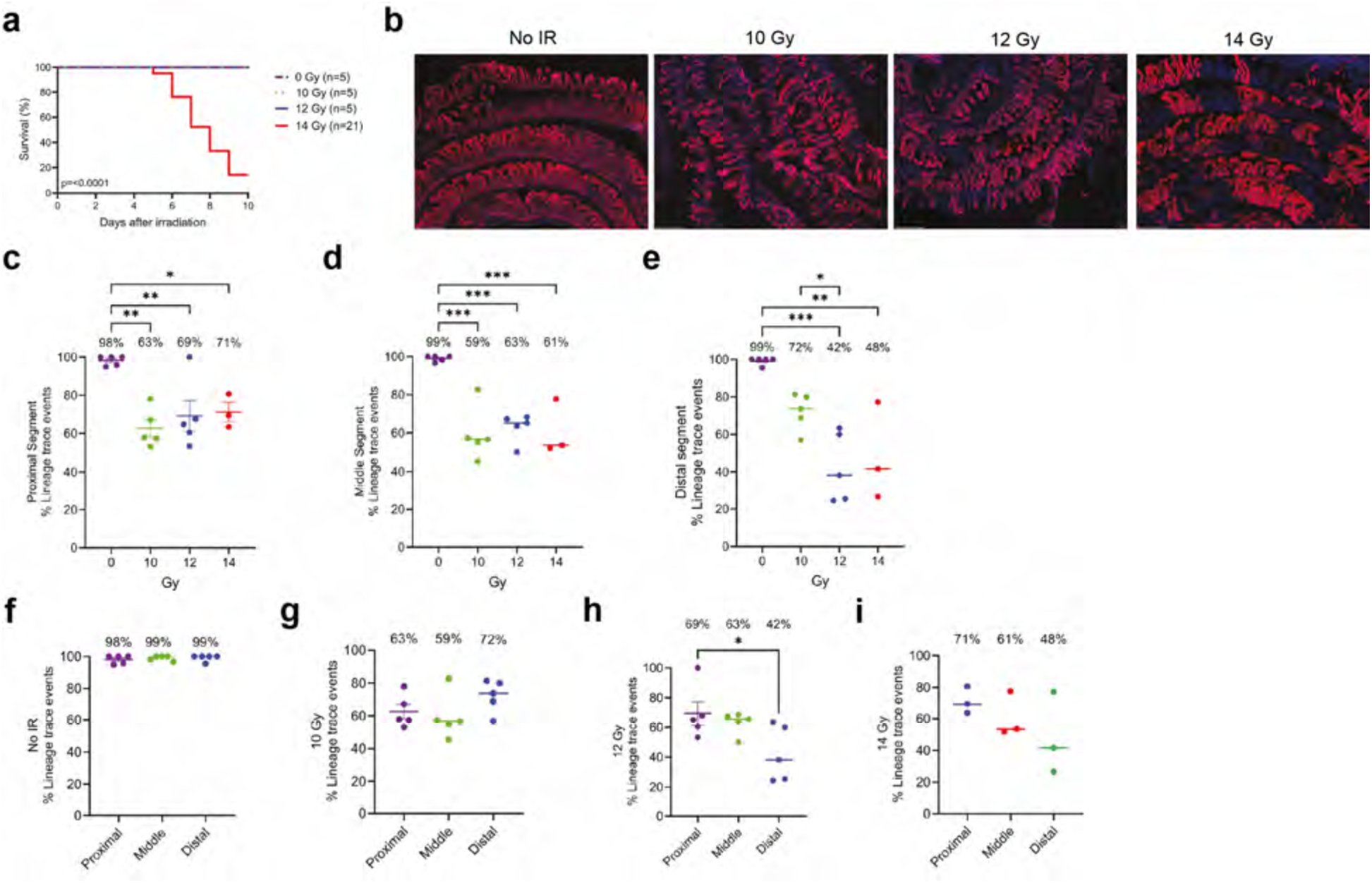
Lineage tracing (2 days pre-IR TAM labeling harvested 10 days post-IR) **(a)** Survival of Lgr5-CreERT2; Rosa26-LSL-tdTomato mice given TAM 2 days prior to 0-14 Gy SBI. Mice were followed for survival for 10 days and survivors’ small intestines were harvested. No IR n=5, 10 Gy SBI n=5, 12 Gy SBI n=5, 14 Gy SBI n=21. p<0.0001 per log-rank test. **(b)** Representative 2.5X images of surviving irradiated Lgr5-CreERT2; Rosa26-LSL-tdTomato small intestines given tamoxifen TAM 2 days prior to harvesting 10 days post-IR. Scale bar 500um. **(c-e)** % lineage trace events per segment (proximal, middle, distal) 0-14 Gy SBI. Each dot represents one mouse. Proximal segment p=0.001 No IR vs. 10 Gy SBI; p=0.007 No IR vs. 12 Gy SBI; p=0.02 No IR vs. 14 Gy SBI. Middle segment p=0.0001 No IR vs. 10 Gy SBI; p=0.0003 No IR vs. 12 Gy SBI; p=0.0008 No IR vs. 14 Gy SBI. Distal segment p=0.0002 No IR vs. 12 Gy SBI; p=0.0019 No IR vs. 14 Gy SBI; p=0.03 10 Gy SBI vs. 12 Gy SBI. All statistics were performed via nonparametric one-way ANOVA. **(f-i)** % lineage trace events per irradiation dose 0-14 Gy SBI/segment. Each dot represents one mouse. 12 Gy SBI proximal vs. distal p=0.04. All statistics were performed via nonparametric one-way ANOVA.

**Extended Data Fig. 8:**
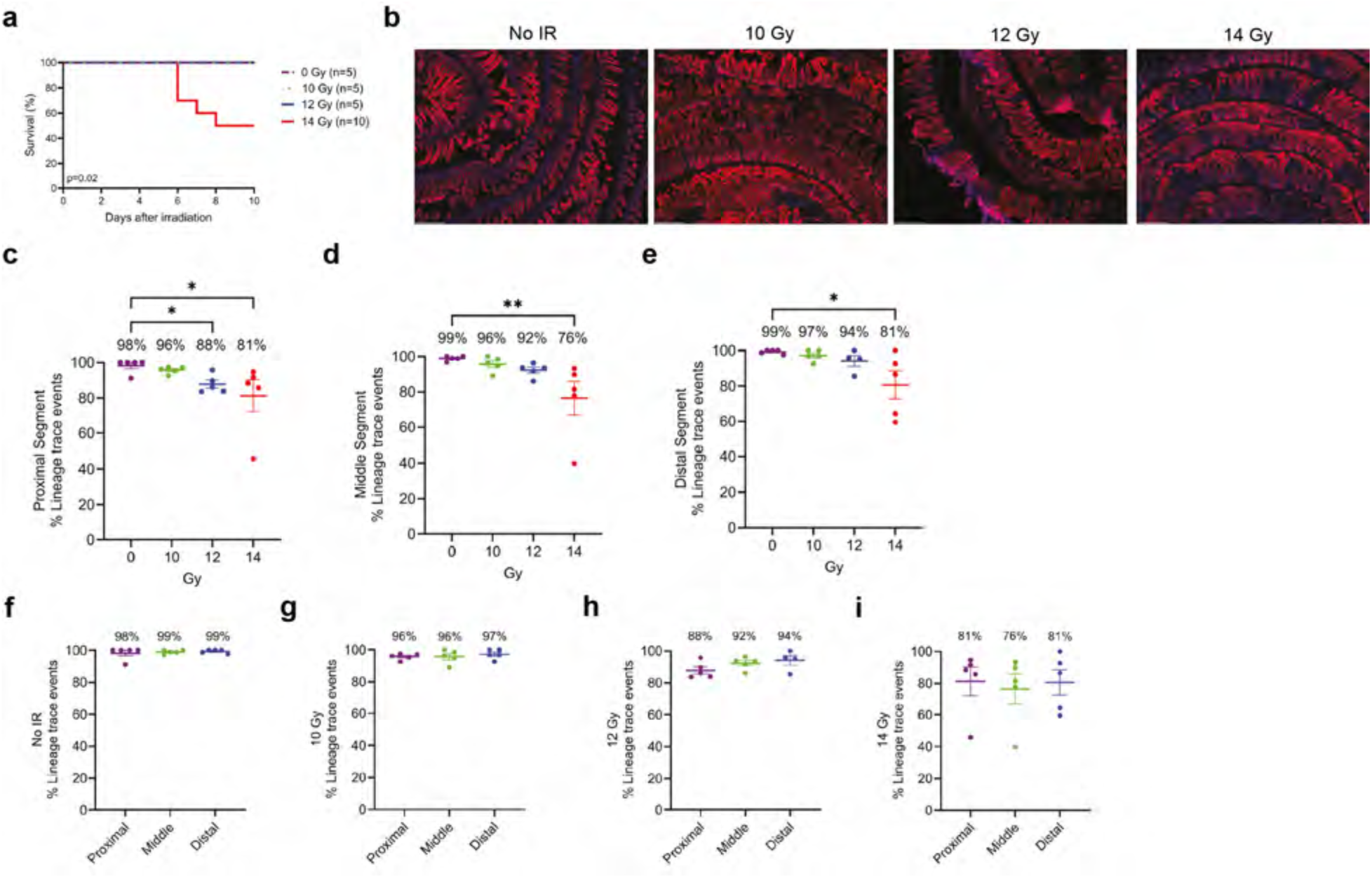
Lineage tracing (7 days pre-IR TAM labeling, harvested 10 days post-IR) **(a)** Survival of Lgr5-CreERT2; Rosa26-LSL-tdTomato mice given TAM 7 days prior to 0-14 Gy SBI. Mice were followed for survival for 10 days and survivors’ small intestines were harvested. No IR n=5, 10 Gy SBI n=5, 12 Gy SBI n=5, 14 Gy SBI n=10. p=0.02 per log-rank test. **(b)** Representative 2.5X images of surviving irradiated Lgr5-CreERT2; Rosa26-LSL-tdTomato small intestines given TAM 7 days prior to harvesting 10 days post-IR. Scale bar 500um. **(c-e)** % lineage trace events per segment (proximal, middle, distal) 0-14 Gy SBI. Each dot represents one mouse. Proximal segment p=0.01 No IR vs. 12 Gy SBI; p=0.01 No IR vs. 14 Gy SBI. Middle segment p=0.001 No IR vs. 14 Gy SBI. Distal segment p=0.04 No IR vs. 14 Gy SBI. All statistics were performed via nonparametric one-way ANOVA. **(f-i)** % lineage trace events per irradiation dose 0-14 Gy SBI/segment. Each dot represents one mouse. All statistics were performed via nonparametric one-way ANOVA.

**Extended Data Fig. 9:**
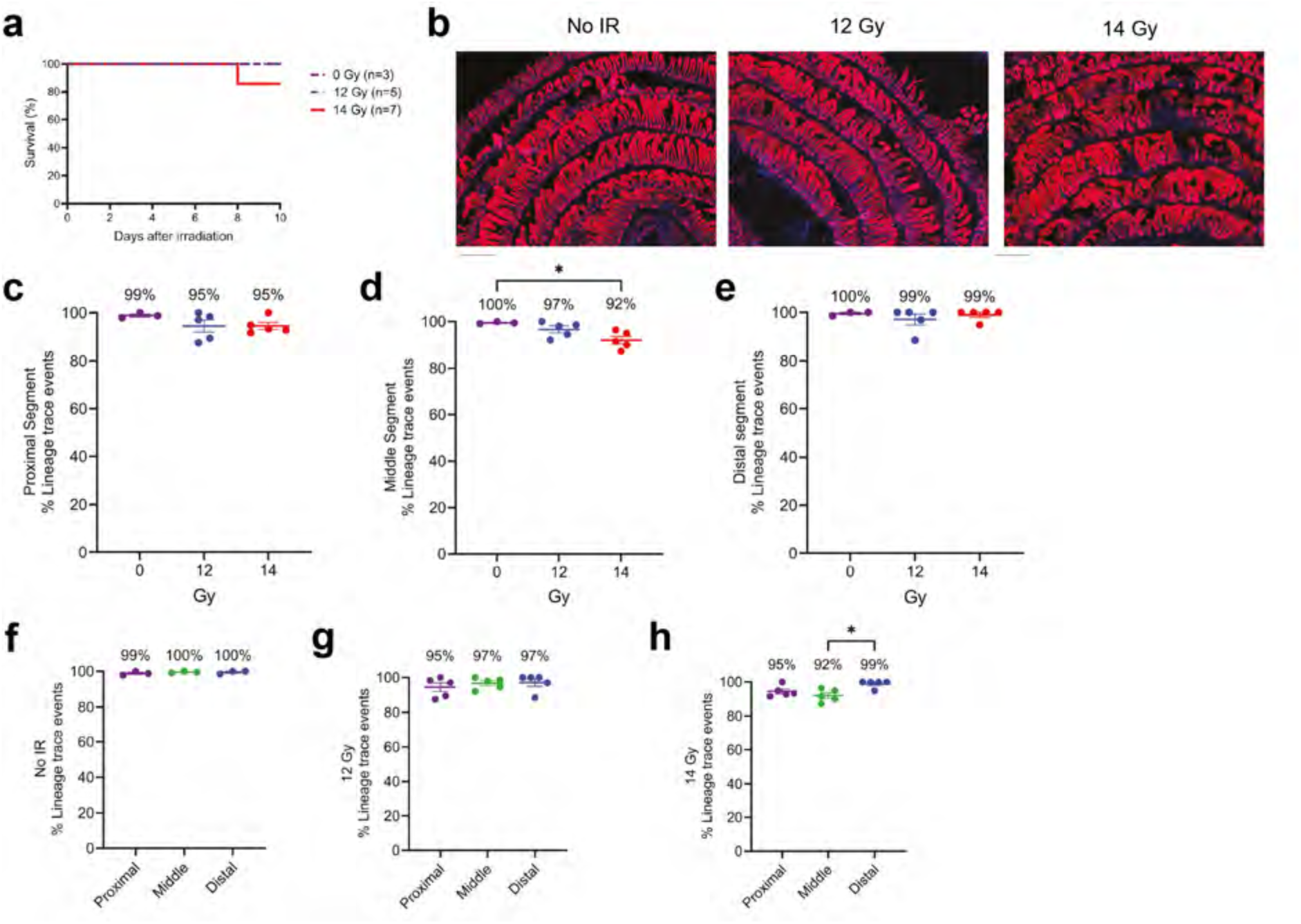
Lineage tracing (31 days pre-IR TAM labeling, harvested 10 days post-IR) **(a)** Survival of Lgr5-CreERT2; Rosa26-LSL-tdTomato mice given TAM 31 days prior to 0, 12, or 14 Gy SBI. Mice were followed for survival for 10 days and survivors’ small intestines were harvested. No IR n=3, 12 Gy SBI n=5, 14 Gy SBI n=7. **(b)** Representative 2.5X images of surviving irradiated Lgr5-CreERT2; Rosa26-LSL-tdTomato small intestines given TAM 31 days prior to harvesting 10 days post-IR. Scale bar 500um. **(c-e)** % lineage trace events per segment (proximal, middle, distal) 0, 12, or 14 Gy SBI. Each dot represents one mouse. Middle segment p=0.02 No IR vs. 14 Gy SBI. All statistics were performed via nonparametric one-way ANOVA. **(f-h)** % lineage trace events of irradiation dose/segment. 14 Gy middle vs. distal p=0.03. Each dot represents one mouse. All statistics were performed via nonparametric one-way ANOVA.

**Extended Data Fig. 10:**
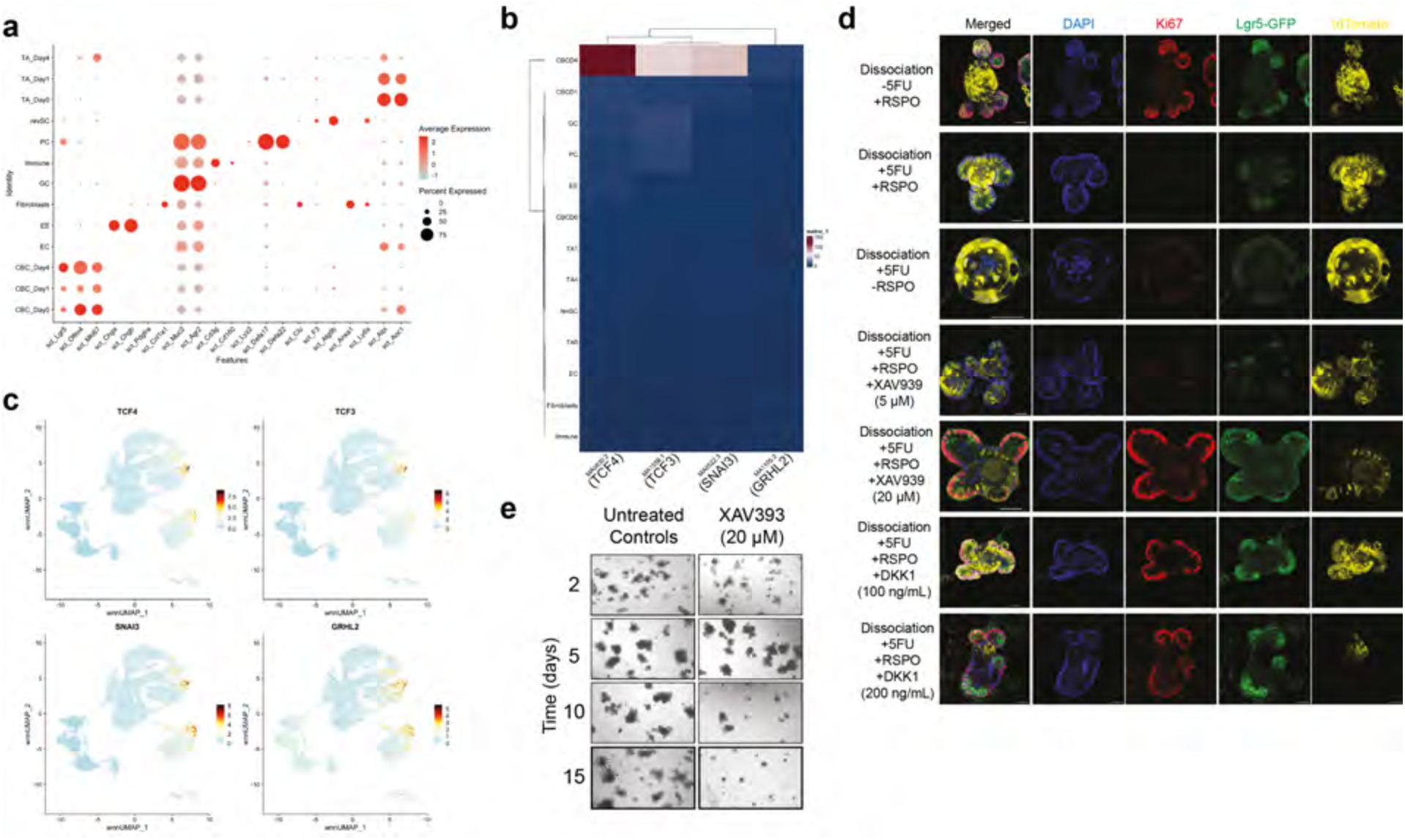
Chromatin accessibility and functional assays of Wnt/B-catenin signaling **(a)** Dot plot of scRNA of multi-omics data depicting expression and percentage of cells for various intestinal cell markers used before. **(b)** Heatmap of TCF4, TCF3, SNAI3, and GRHL2 motif expression in all cell type clusters in WNN dataset. Accessibility of motif TCF4, TCF3, SNAI3, and GRHL2 for WNN dataset. **(c)** Feature plot of expression of TCF4, TCF3, SNAI3, and GRHL2 in WNN dataset. **(d)** Images of Lgr5DTR-EGFP and Clu^CreERT^^2^, Rosa26^loxp-STOP-loxp-tdTomato^ organoids with treatment combinations with 5-FU, RSPO, XAV939 and DKK1 and various concentrations of inhibitors. Organoids were stained with Dapi and Ki67. **(e)** c57 mouse intestinal organoids over the span of 15 days, untreated and treated with XAV393.

**Supplementary Table 1: Metadata corresponding to cells in reference dataset for cell identity, phase, beta-catenin signature and replicating signatures.**

**Supplementary Table 2: Differential gene expression analysis for all cell types in the reference dataset.**

**Supplementary Table 3: Differential gene expression analysis for CBC1 and CBC2 in the reference dataset.**

**Supplementary Table 4: Differential gene expression analysis for all cell types in multiomics dataset.**

**Supplementary Table 5: Motif enrichment analysis for all cell types in the multiomics dataset.**

